# A cooperative response to endocardial NOTCH signaling stimulation regulates transcriptional activity during cardiac valve development and disease

**DOI:** 10.1101/2023.03.23.533882

**Authors:** Luis Luna-Zurita, Brenda Giselle Flores-Garza, Dimitrios Grivas, José Luis de la Pompa

## Abstract

**Background:** The endocardium is a crucial signaling center for cardiac valve development and maturation. Genetic analysis has identified several human endocardial genes whose inactivation leads to bicuspid aortic valve (BAV) formation and/or calcific aortic valve disease (CAVD), but knowledge is very limited about the role played in valve development and disease by non-coding endocardial regulatory regions and upstream factors.

**Methods:** We manipulated the NOTCH signaling pathway in mouse embryonic endocardial cells by short-term and long-term co-culture with OP9 stromal cells expressing NOTCH ligands and treatment with the γ-secretase inhibitor RO4929097, defining the transcriptional profile associated to each condition. The endocardial chromatin accessibility landscape for each condition was defined by high-throughput sequencing (ATAC-seq) determination of transposase-accessible chromatin. In vitro and in vivo models carrying deletions of different non-coding regulatory elements were generated by CRISPR-Cas9 gene editing.

**Results:** We identified primary and secondary transcriptional responses to NOTCH ligands in the mouse embryonic endocardium. By integrating our gene expression data with data from developing valves of mice with NOTCH loss-of-function and from human valve calcification samples, we were able to identify a NOTCH-dependent transcriptional signature in valve development and disease. Further, by defining the endocardial chromatin accessibility landscape after NOTCH pathway manipulation and integrating with in vivo data from developing mouse endocardium and adult human valves, we were able to identify a set of potential non-coding regulatory elements, validate representative candidates, propose co-factors interacting with them, and define the timeframe of their regulatory activity. Analysis of the transcriptional repression driven by NOTCH activation revealed cooperation between the NOTCH and HIPPO pathways in the endocardium during cardiac valve development.

**Conclusions:** Transcriptional regulation in the embryonic endocardium after NOTCH pathway stimulation occurs in a sequential manner and requires the participation of several factors. NOTCH not only triggers the transcriptional activity of the non-coding elements recognized by these factors, but also represses those elements whose activity negatively affects the development and homeostasis of the cardiac valves.

**Novelty and Significance:** WHAT IS KNOWN?

- The embryonic endocardium is a source of cell types and a crucial signaling center for cardiac valve development and maturation.
- Endocardial NOTCH pathway activity patterns specific cellular behaviors that will give rise to the cardiac valve primordia, guide their maturation, and maintain adult valve homeostasis.
- NOTCH signaling abrogation is associated with human valve dysmorphology (bicuspid aortic valve) and adult calcific aortic valve disease (CAVD).

WHAT NEW INFORMATION DOES THIS ARTICLE CONTRIBUTE?

- The existence of a primary transcriptional response after short-term NOTCH activation that is enhanced after sustained ligand stimulation, and a secondary response triggered after long-term NOTCH activation.
- We have identified a NOTCH-dependent transcriptional signature specific for valve development and disease.
- We have defined the endocardial chromatin accessibility landscape after NOTCH pathway manipulation, and proposed a set of potential cofactors and the time frame of their activity involved in the NOTCH-dependent transcriptional response.
- We have identified a set of potential non-coding regulatory elements involved in valve development and disease, and evaluated them in vitro and in vivo.
- We have identified the cooperation between the NOTCH and HIPPO pathways in the endocardium during cardiac valve development.

## Introduction

The endocardium is a specialized endothelium covering the inner surface of the myocardial wall^1^. During cardiac development, the embryonic endocardium is a source of cell types and signals that are essential for the generation of a functional heart ^2^. One key signaling pathway for these developmental processes is initiated by the membrane receptor Notch. Delta and Jagged/Serrate ligands expressed on neighboring cells trigger a series of cleavage events in the Notch receptors (Notch1-4) that ultimately result in the generation of the NOTCH intracellular domain (NICD) through γ-secretase activity ^3^. The liberated NICD translocates to the nucleus, where it binds to a preexisting transcriptional complex including RBPJ to induce target gene expression ^4^. At early stages of cardiac development, the endocardium and myocardium are separated by an extracellular matrix (ECM), the cardiac jelly. This structure is later progressively reabsorbed in the chambers, but in the presumptive valve territories it is colonized by mesenchymal cells derived from endocardial-to-mesenchymal transition (EMT). The resulting endocardial cushions form the valve primordia and are progressively remodeled through spatiotemporally regulated proliferation and apoptosis of valve interstitial cells together with tightly controlled synthesis and organization of the ECM ^5^. EMT of the early endocardium in the atrioventricular canal (AVC) and outflow tract (OFT) regions requires the sequential activation of Notch by Delta4 (Dll4). Jagged1 (Jag1) signaling is dispensable for this process, but the later remodeling of the valve primordia requires induction of Notch activity by Jag1 expressed on endocardial cells ^6^. Dysregulation of NOTCH signaling is associated with human valvular congenital heart disease (CHD) ^7–9,10, 11^ and acquired calcific aortic valve disease (CAVD) ^12–14^.

Much recent effort has been directed at the discovery of distal non-coding regulatory genomic regions owing to their potential implication in organ development and disease. Particular emphasis has been placed on the use of genome-wide association studies (GWAS) to identify distally located disease-risk variants, especially those associated with valve disease ^15–19^. These studies highlight the importance of identifying regulatory elements affected by these variants, since the large size of the non-coding genome relative to coding sequences makes it an important mutational target with potential disease-causing consequences. Despite the pivotal role of NOTCH signaling in valve development and disease, the NOTCH-dependent transcriptional program and regulatory features in the embryonic endocardium remain unknown. There is thus a clear need to identify NOTCH-activity-dependent regulatory elements, their interacting factors, and the dynamics of their activity.

Here, by manipulating the Notch signaling pathway in mouse embryonic endocardial cells (MEEC), we identify a complex and sequential transcriptional response in which short-term and long-term Notch stimulation produces different outputs, and we describe the chromatin accessibility dynamics responsible for each response. Integration of these findings with transcriptional and epigenomic data from developing and adult valves allowed us to identify a Notch-dependent transcriptional signature in valve development and disease and a new set of non-coding regulatory elements potentially regulated by multiple co-factors. Although the response to short-term stimulation was stronger for Jag1, we found no ligand-specificity in the chromatin and transcriptional output driven by Jag1 or Dll4. An analysis of the transcriptional repression driven by Notch activation exposed antagonistic roles of Notch and Yap/Taz in the endocardium during cardiac-valve development.

## Methods

Full methods are provided in the Online Data Supplement. Animal studies were approved by the CNIC Animal Experimentation Ethics Committee and by the Community of Madrid (Ref. PROEX 155.7/20). All animal procedures conformed to EU Directive 2010/63EU and Recommendation 2007/526/EC regarding the protection of animals used for experimental and other scientific purposes, enacted in Spanish law under Real Decreto 1201/2005.

## Results

### Jag1 and Dll4 ligands differ in efficiency but have similar gene targets in mouse embryonic endocardial cells

Endocardial Notch activity in the presumptive valve territory is sequentially regulated by Dll4 and Jag1. At E9.5, Notch1 activity is found in the presumptive valve endocardium, where the ligands Dll4 and Jag1 are both expressed^6^; however, at this stage only the lack of *Dll4* affects endocardial cushion tissue cellularization (Figure S1A)^6^. In contrast, during valve primordium remodeling, Jag1 was the only ligand detected in the valve region, in the myocardium and endocardium^6^. The requirement of Jag1-Notch1 signaling for valve morphogenesis is supported by the observation that abrogation of endocardial Jag1 signaling leads to bicuspid aortic valve (BAV) (Figure S1B)^6^. To define the transcriptional effect of Dll4- and Jag1-induced Notch signaling on the embryonic endocardium, we cultured mouse embryonic endocardial cells (MEEC) ^6, 20, 21^ over stromal OP9 cells expressing Dll4 or Jag1 for 6 or 24 hours. Short-term (6h) stimulation with Dll4 yielded only 47 differentially expressed genes (DEGs), contrasting with 258 DEGs induced by 6h stimulation with Jag1 (Figure 1A, top, Table S1). Long-term (24h) stimulation significantly increased the number of DEGs for both ligands, to 516 with Dll4 and 1,118 with Jag1 (Figure 1A, down, Table S1). Despite the between-ligand differences in the number of DEGs, the gene expression changes showed very good correlation (Figure 1B, S1E), with most DEGs induced by Dll4 also induced by Jag1, and the main difference being the intensity and not the direction of the fold changes (Figure 1C, S1C,D). These results show that Jag1 is the more efficient ligand in terms of signaling output and reveal a lack of ligand-specific target genes in the embryonic endocardium.

**Figure 1.**
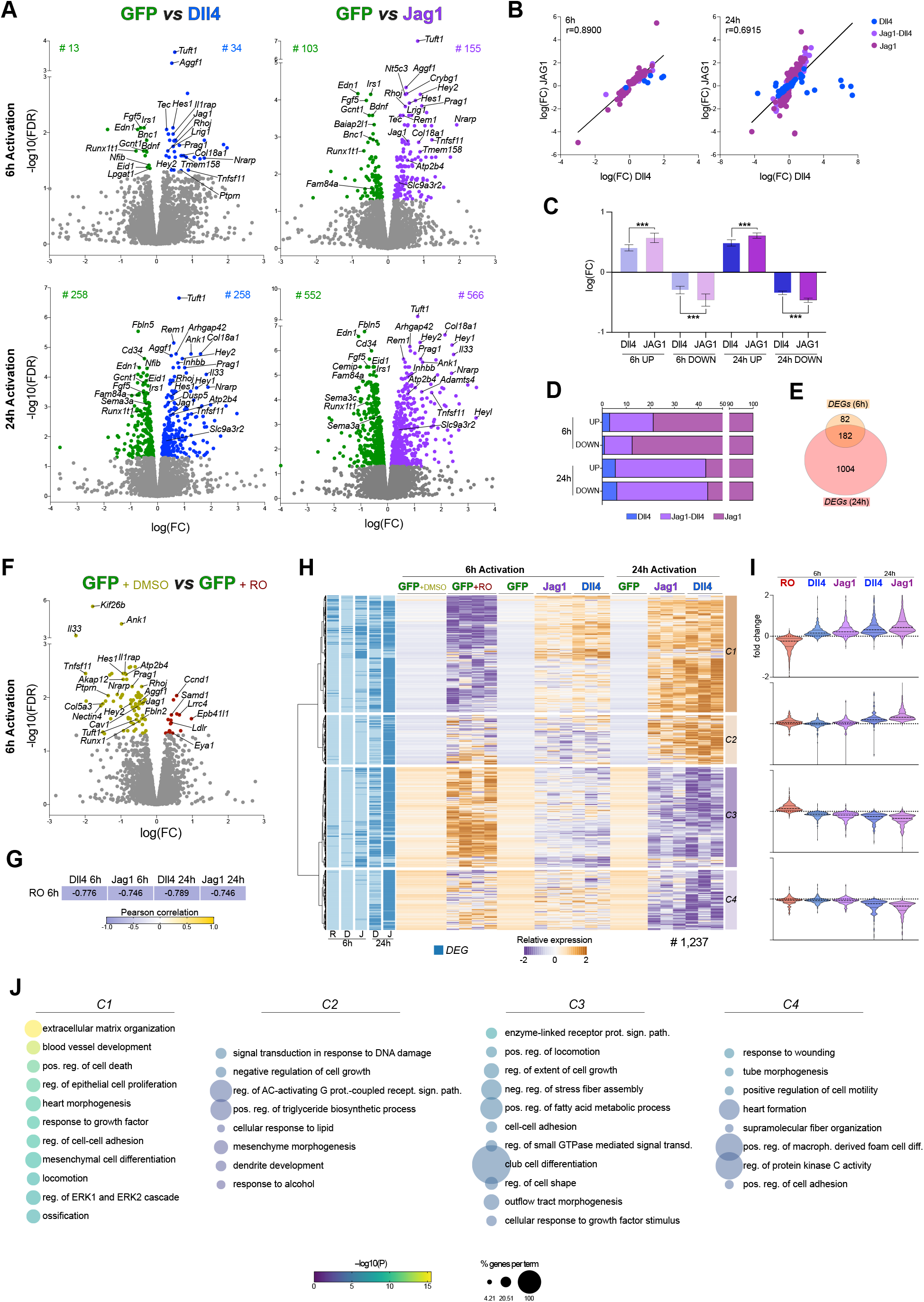
Transcriptional analysis in mouse embryonic endocardial cells after the manipulation of Notch-pathway activity. (A) Volcano plot showing fold changes in expression and associated -log^10^(FDR) values for each detected gene altered in mouse embryonic endocardial cells (MEEC) upon stimulation with Notch ligands expressed on OP9 cells. MEEC were co-cultured over OP9 cells expressing GFP (control), Dll4, or Jag1 for 6 (top) or 24 hours (bottom). Differential gene expression versus co-culture with OP9-GFP cells (FDR < 0.05) is indicated in blue (upregulation with OP9-Dll4), purple (upregulation with OP9-Jag1), and green (downregulation). (B) Scatter plot comparing fold expression changes after co-culture with OP9-Dll4 versus OP9-Jag1 for each DEG detected after 6 (left) or 24h (right). (C) Bar graph showing mean fold changes in genes upregulated and downregulated after Dll4 and Jag1 stimulation for 6h (short-term; 6h) and 24 h (long-term; 24h). (D) Stacked bar plot showing the percentage of differentially expressed genes dependent on Jag1 and/or Dll4 after 6 or 24h stimulation. (E) Venn diagram representing the proportion of DEGs detected after 6 or 24h stimulation. (F) Volcano plot showing fold changes in expression and associated -log^10^(FDR) values for each detected gene altered in MEEC co-cultured with OP9-GFP cells + RO4929097 (RO) for 6h. Differential gene expression with RO versus DMSO (FDR < 0.05) is indicated in red (upregulation) and green (downregulation). (G) In the DEGs detected in (F), Pearson correlation between changes in gene expression due to RO activity and the changes due to Dll4 and Jag1 activation for 6 or 24h. (H) Hierarchical clustering of genes differentially expressed in MEEC treated with RO or with OP9-expressed Dll4 and Jag1. (I) Violin plots representing the fold expression changes of genes detected in each cluster. (J) Enriched GO (biological function) terms for each cluster. The size of the nodes represents the percentage of genes in the cluster associated with the term. The color code represents the -log^10^(p value) for the enrichment.

### Mouse embryonic endothelial cells initiate a sequential transcriptional response after Notch ligand activation by neighboring cells

To determine if the detected DEGs are altered as a direct (primary) or downstream effect of the transcriptional response mediated by the NICD-RBPJ complex, we co-cultured MEEC over control OP9 for 6 hours in the presence of DMSO (vehicle) or the NOTCH-signaling inhibitor RO4929097 (RO, see Methods) (Figure 1F, Table S1). Changes in gene expression induced by RO inversely correlated with the changes triggered by Dll4 and Jag1 signaling (Figure 1G), confirming the crucial role of NICD-RBPJ in the detected transcriptional alterations. The RO-induced changes also allowed us to define baseline gene expression for comparison with the changes observed after stimulation with Dll4 or Jag1. Comparison of the transcriptional profiles obtained in the three experimental conditions (6h and 24h ligand stimulation and RO administration) revealed a two-phase response to Notch ligand exposure: a primary transcriptional response after NICD nuclear translocation (clusters C1 and C3) that was sustained and enhanced over time; and a secondary response (clusters C2 and C4), triggered after long-term ligand exposure (Figure 1H-I, Table S1). In clusters C1 and C3, RO-mediated inhibition of Notch receptor cleavage reduced (C1) or increased (C3) gene expression levels, whereas expression gradually increased (C1) or declined (C3) during continuous exposure to Notch ligands (Figure 1H-I). In contrast, the transcriptional response in clusters C2 and C4 was observed only after 24 hours of ligand exposure (Figure 1H-I). Biological function terms overrepresented in a Gene Ontology (GO) analysis of the primary transcriptional response (C1, C3) included extracellular matrix, cell-cell adhesion, locomotion, stress fiber assembly, regulation of the ERK1 and ERK2 cascades, and GTPase-mediated signal transduction (Figure 1J, Table S1). C2 genes were mostly associated with transcriptional activity and metabolism, whereas function terms overrepresented in C4 included response to wounding, tube morphogenesis, cell motility, and macrophage differentiation (Figure 1J, Table S1). The three experimental conditions thus revealed a complex transcriptional response to NOTCH ligands. A set of genes is activated after short-term ligand stimulation, as a direct response to NOTCH activation; expression of these genes is maintained and enhanced after sustained ligand stimulation. Long-term exposure to Notch ligands triggers a later transcriptional response of genes unresponsive to short periods of Notch pathway activity.

### A Notch-dependent gene expression signature associated with cardiac valve development and disease

To assess whether the transcriptional responses identified in MEEC also occur in intact embryonic endocardium in vivo, we examined two available transcriptional datasets: one from a mouse model of heart valve development, and another from human heart valves in health and disease. The mouse dataset is from the semilunar valves of *Jag1^flox/flox^;Nkx2-5-Cre* mice, which lack Jag1 in the myocardium and endocardium and are characterized by hyperplastic and dysmorphic valves at E15.5 and a high rate of BAV ^6^. Analysis of this dataset showed that the set of highly expressed genes in E14.5 WT micro-dissected semilunar valves is enriched in DEGs from clusters C1 and C2 (increased expression upon Notch ligand stimulation) (Figure 2A, left). In contrast, the semilunar valves of *Jag1^flox/flox^;Nkx2-5-Cre* mice show overexpression of many of the DEGs repressed by Notch activity (clusters C3 and C4) (Figure 2A, left). These findings confirm that the transcriptional response to Notch signaling observed in vitro recapitulates the natural process operating in developing semilunar valves.

**Figure 2.**
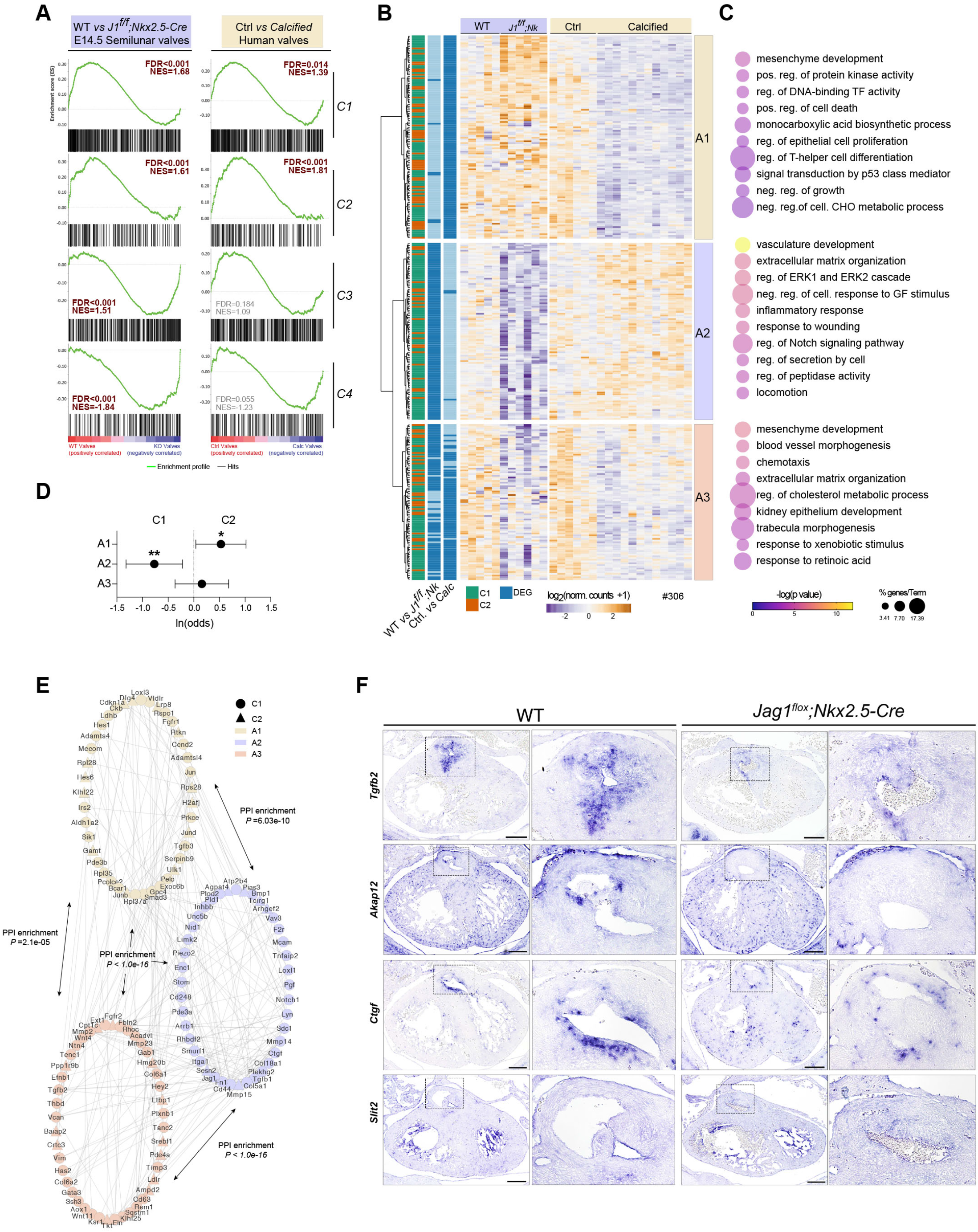
The Notch-dependent transcriptional response in mouse embryonic endocardial cells operates in mouse valve development and human valve disease. (A) Gene-set enrichment analysis showing the distribution of DEGs located in clusters C1-C4 from Figure 1 H-J within the ranking of all genes identified in E14.5 WT and *Jag1^flox^;Nkx2.5-Cre* semilunar valves (left) and in control (Ctrl) and calcified adult human valves (right). FDR values represent the statistical significance of the enrichment score. NES, normalized enrichment score. (B) Hierarchical clustering of cluster C1 and C2 genes significantly enriched in E14.5 WT versus *Jag1^flox^;Nkx2.5-Cre* semilunar valves or in Ctrl versus calcified human valves. (C) Enriched GO (biological function) terms for each cluster. The size of the nodes represents the percentage of genes in the cluster associated with the term. The color code represents the -log^10^(p value) for the enrichment. (D) Odds ratios for finding C1, C2, or C3 DEGs in clusters A1, A2, or A3. (E) Protein-protein interactome prediction for genes identified in clusters A1, A2, or A3. The number of predicted interactions between clusters was statistically significant. (F) ISH analysis of *Tgfb2*, *Akap12*, *Ctgf*, and *Slit2* expression in heart sections from E14.5 WT and *Jag1^flox^;Nkx2.5-Cre* embryos. Magnified views of the aortic valve are shown to the right.

Another recent transcriptional analysis identified an immune-metabolic gene expression signature in human adult calcified bicuspid valves (cBAV) and tricuspid valves (cTAV) ^22^. Aortic-valve calcification is the most common form of valvular heart disease and is repressed by NOTCH activity ^10, 13, 23^. Consistent with this proposed role of NOTCH activity, analysis of this dataset showed that C1 and especially C2 DEGs were significantly more abundant in control adult valves, indicating downregulation of these genes in CAVD (Figure 2A, right). In contrast, genes repressed by NOTCH ligands showed no significant difference between samples (Figure 2A, right). This observation indicates that transcriptional changes associated with human valve calcification involve the repression of genes positively regulated by Notch activity in the endocardium.

Given the overrepresentation of clusters C1 and C2 (positive primary and secondary transcriptional responses to Notch activity) in the controls of both these models versus the mutant or disease state, we investigated the specific role of these clusters in each setting. Hierarchical clustering of gene expression changes revealed the downregulation of genes specifically in human calcified aortic valves (A1), in developing mouse valves lacking Jag1 signaling (A2), and in both settings (A3) (Figure 2B). GO analysis revealed a pronounced involvement of genes related to the endothelium, extracellular matrix, and ERK signaling in cluster A2 (downregulated in *Jag1^flox/flox^;Nkx2.5-Cre* semilunar valves) (Figure 2C), functions similar to those associated with cluster C1 in the in vitro model (Figure 1J). Cluster A1 (human calcified valves) was associated with metabolic and proliferative functions (Figure 2C), comparable to cluster C2 (Figure 1J). Consistent with this finding, C1 DEGs were overrepresented in cluster A2, whereas C2 DEGs were more represented in cluster A1 (Figure 2D), suggesting that the secondary transcriptional response to Notch activity plays an important role in preventing aortic valve calcification. Despite the observed differences, a predicted protein-protein interaction network constructed from the DEGs identified in clusters A1-3 revealed significant interactivity both within each cluster and among the three of them, revealing close association and common functions among the genes detected in vivo (Figure 2E). Further assessment of the effect Jag1-Notch1 signaling blockade in developing mouse aortic valves identified reduced expression of specific genes in clusters A1 (*Akap12*), A2 (*Ctgf*), and A3 (*Tgfb2* and *Slit2*) (Figure 2F). Together, these results confirm the relevance of our in vitro dataset to the study of valve development and disease.

### Endocardial secretion of Notch-dependent soluble proteins

Endocardial Notch signaling involves the expression and secretion of soluble factors that modify adjacent ECM and/or signal to nearby myocardium (D’Amato et al., 2016),^21, 24^. We previously described the Notch-dependent endocardial secretome after stimulation of MEEC with recombinant NOTCH ligands ^21^. Here, we performed RNA-seq on MEEC stimulated with recombinant ligands (see Methods), finding good correlation with the transcriptional changes elicited by NOTCH ligand-receptor interaction in the MEEC–OP9 co-culture assay (Figure S2A, Table S3). The consistency between these models allowed us to integrate our transcriptional dataset with the Notch-dependent secretome, identifying gene sets whose endocardial expression and protein secretion change upon Notch manipulation in the same way (clusters II and IV) or in the opposite way (clusters I and III) (Figure S2B, Table S3). Integration of these analyses with the in vivo data described in Figure 2A identified Tgfb2, Fbln2, Col6a1, and Timp3 as proteins whose secretion and expression in valve endocardium is affected by Notch abrogation during valve development in mice (Figure S2C, Table S3) and in human CAVD (Figure S2D, Table S3). These observations highlight the value of our dataset for understanding the importance of NOTCH signaling in valvulogenesis.

### Notch-pathway manipulation in mouse embryonic endocardial cells alters chromatin accessibility

To identify the regulatory features orchestrating the Notch-dependent transcriptional cascade in the endocardium, we used ATAC-seq to study changes in chromatin accessibility in MEEC after OP9 co-culture (OP9-Dll4 and OP9-Jag1 cells for 6 and 24 hours and OP9 + RO for 6 hours). Using relaxed selection parameters (see Methods), we defined 4,229 ATAC-seq peaks as differential accessibility regions (DARs, 1,483 after 6h stimulation, 2,690 after 24h, and 1,092 after RO treatment; Figure 3A-B, Table S4), finding reduced and increased accessibility in each experimental condition (Figure 3B, Table S4). Whereas the gene expression analysis revealed that the same genes were affected inversely by Notch inhibition vs activation (Figure 1G, S1C), changes in the chromatin accessibility profile were highly specific for each condition (inhibition and short-term and long-term activation) (Figure S3A-B), with very few DARs occurring in more than one experimental condition (Figure 3A). This specificity for each condition was well illustrated after clustering the 4,229 DARs according to their accessibility changes upon Notch pathway activation or inhibition (Figure S3A). Each cluster (a-g) was characterized by a major accessibility change in one of the experimental conditions, and no general pattern was evident (Figure S3A).

**Figure 3.**
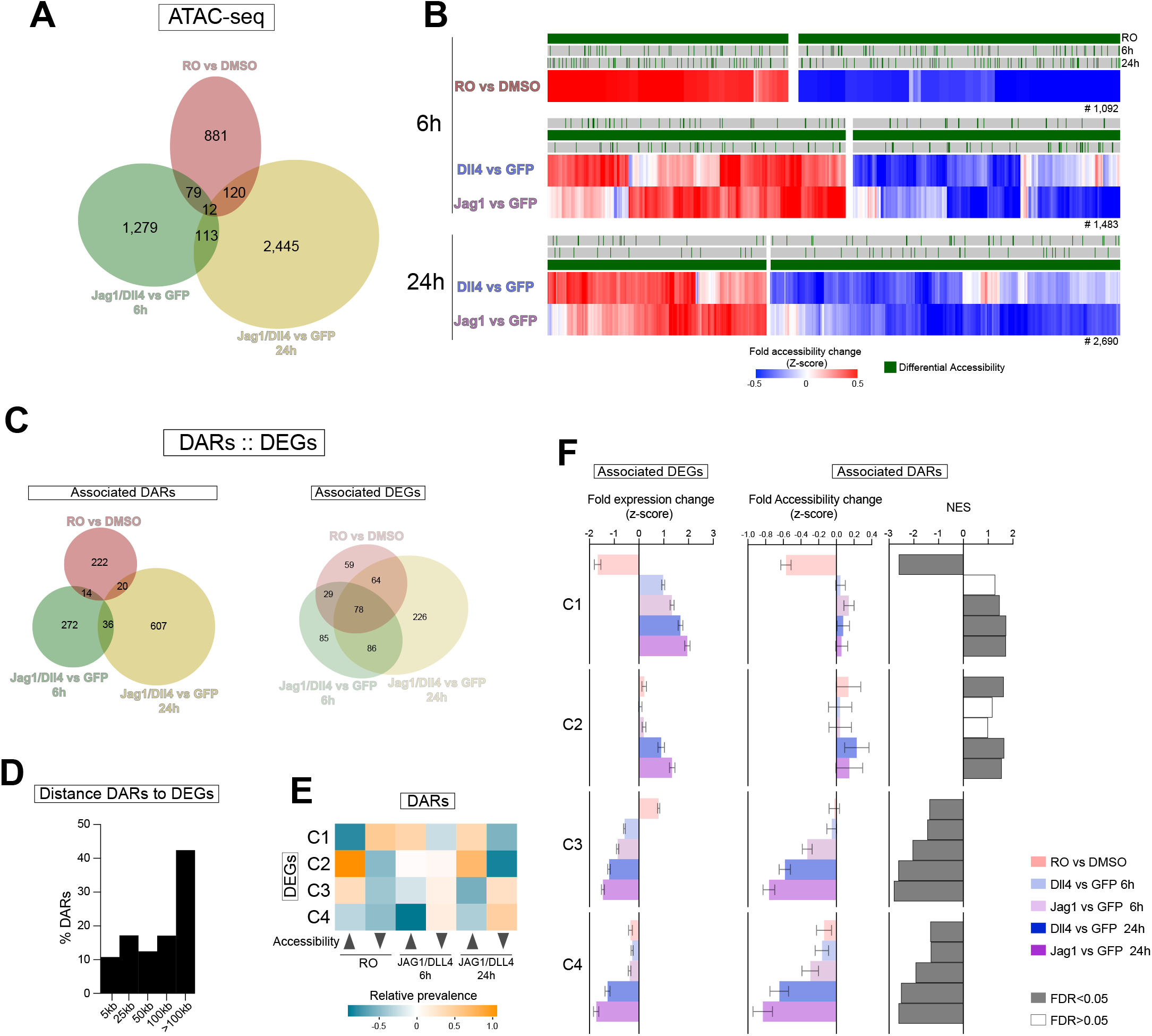
Chromatin accessibility changes and association with gene expression in mouse embryonic endocardial cells after the manipulation of Notch pathway activity. (A) Left. Overlap between differential accessibility regions (DARs) detected after RO treatment for 6h and Notch-ligand stimulation for 6h and 24h. (B) Hierarchical clustering of DARs in each experimental condition. (C) For DAR–DEG pairs, overlap between DAR peaks (left) and DEGs (right) detected after RO treatment for 6h and Notch-ligand stimulation for 6h and 24h. (D) Distance of DARs from the TSS of the associated DEG. (E) Relative prevalence of chromatin accessibility patterns in the regulatory domains of DEGs from each expression cluster. (F) Left: Median fold expression change in each experimental condition in DEGs from each expression cluster (C1-C4) associated with any DAR. Center: Median fold accessibility change (z-score) in each experimental condition in DARs associated with DEGs from each expression cluster. Right: GSEA normalized enrichment score (NES) for DARs associated with DEGs in each expression cluster. Error bars, s.e.m.

For example, clusters b and e were characterized by a strongly reduced or increased accessibility after RO treatment, but there was no clear pattern after 6 or 24 hours of ligand exposure. To assess the relevance of each chromatin accessibility profile to the Notch-dependent transcriptional response, we defined a regulatory domain for each DEG (using GREAT, see Methods) and identified the DARs overlapping with each regulatory domain. Using this approach, we defined 1,252 DAR–DEG pairs (Table S4). Contrasting the limited overlap among DARs between conditions (Figure 3C left), DEGs in these pairs were frequently associated with more than one DAR, each one detected in different experimental conditions (Figure 3C right). DARs showed a genome-wide distribution, with 81.7% detected in intergenic or intragenic regions (Figure S3C). Consistent with this distribution, DARs were preferentially located in regions distal to the TSS of their associated DEGs, with 90% of them at a distance of more than 5kb (Figure 3D). Despite this distal location, changes in accessibility and gene expression in DAR–DEG pairs showed good correlation (Figure S3C), supporting a potential regulatory role of the DARs over their paired DEGs.

To identify specific changes in accessibility associated with expression changes of the putative DEGs, we assessed the prevalence of each accessibility pattern in the regulatory domains of DEGs in each expression cluster (C1-C4) (Figure 3E). Consistent with the expression pattern of C1 DEGs (Figure 1H, 3F left), the most prevalent chromatin accessibility pattern in the DARs associated with C1 DEGs featured repression after RO exposure and increased accessibility after ligand activation (Figure 3E, F center and right). In the regulatory domains associated with cluster C2 (the late positive response to ligand activation; Figure 1H, 3F left), the most prevalent chromatin pattern was increased accessibility after long-term ligand stimulation (Figure 3E, F center and right). These results suggest that the induction and maintenance of the expression of C1 DEGs requires the sequential participation of distinct genomic elements, whereas for C2 DEGs gene expression appears to depend on regions responding to long-term ligand activity.

Similarly, the early and late transcriptional repression in clusters C3 and C4 (Figure 1H, 3F left) was reflected in a general repression of chromatin accessibility in the associated regulatory regions after Notch ligand stimulation (Figure 3E, F). Interestingly, for the response to RO, consistency between changes in DARs and their associated DEGs was only found only for cluster C1, indicating that a major role of the NICD-RBPJ complex is restricted to this cluster. Together, these results suggest complex but well-coordinated transcriptional regulation in response to Notch ligand signaling.

### Multiple transcription factors are involved in the Notch-dependent endocardial transcriptional response

To identify factors interacting with DARs, and thus potentially orchestrating the observed transcriptional changes, we performed a known-motif enrichment analysis (using Homer, see Methods) of DARs showing increased or reduced accessibility in response to each experimental condition. These accessibility patterns were grouped according to whether Notch pathway activation increased accessibility (repressed accessibility after RO treatment and increased accessibility after Notch-ligand stimulation for 6h or 24h; Figure 4A, top three rows) or repressed it (increased accessibility after RO treatment and reduced accessibility after Notch-ligand stimulation; Figure 4A, bottom three rows).

**Figure 4.**
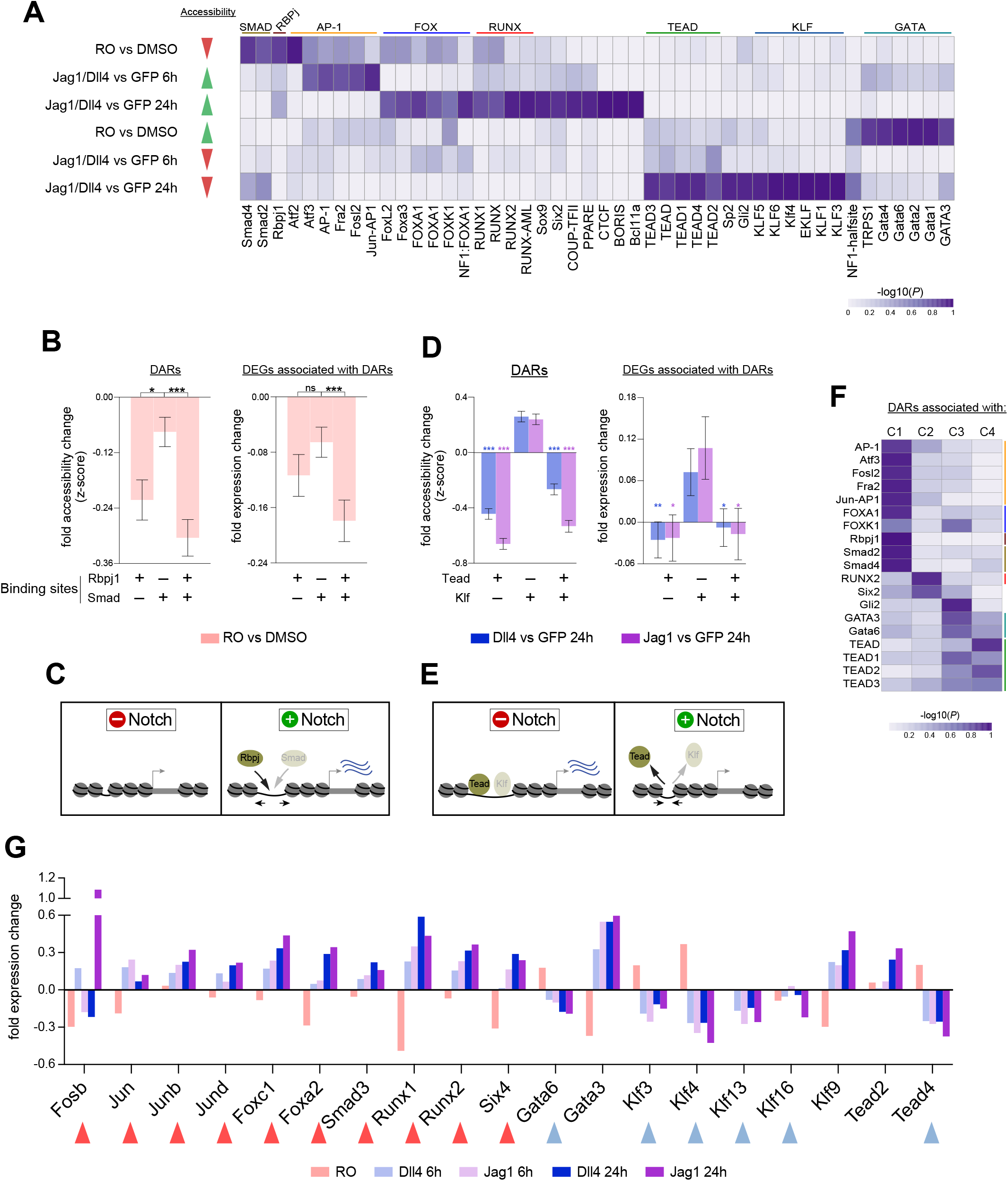
Potential transcription factors controlling gene expression changes in mouse embryonic endocardial cells in response to manipulation of the Notch pathway activity. (A) Heatmap showing the normalized -log10(pvalue) of known binding site motifs detected in at least one of the clusters from Figure 3B. (B) Left. Median fold accessibility change (z-score) after RO treatment in DARs affected by RO and containing RBPj, SMAD, or both binding sites; Right. Median fold expression change after RO treatment in DEGs associated with DARs containing RBPj, SMAD, or both binding sites. (C) Proposed Notch-induced transcription factor occupancy in genomic elements repressed by RO treatment. (D) Left. Median fold accessibility change (z-score) after Dll4/Jag1 stimulation for 24 h in DARs containing TEAD, KLF, or both binding sites. Right. Median fold expression change after Dll4/Jag1 stimulation for 24 h in DEGs associated with DARs containing TEAD, KLF, or both binding sites. (E) Proposed Notch-induced transcription factor occupancy in genomic elements repressed by long term Notch-ligand activity. (F) Heatmap showing the normalized -log10(pvalue) of known binding motifs detected in (A) and overrepresented in DARs associated with DEGs in each expression cluster (C1-C4). (G) Fold expression changes of differentially expressed TFs that have their motif enriched in each DAR cluster. Error bars, s.e.m. In B and D, * p<0.05, ** p<0.01, *** p<0.001 (Mann Whitney test).

RO-driven repression of accessibility was characterized by the presence of binding sites for RBPJ and also for SMAD (Figure 4A, Table S4). NICD-RBPJ forms part of a large transcriptional complex that interacts with specific regulatory elements able to induce the expression of direct Notch target genes ^25^. These specific NICD-RBPJ interacting regulatory elements are likely to occur in those genomic elements whose accessibility was repressed in MEEC after RO treatment in the co-culture Notch-signaling model. Previous studies have reported crosstalk between Notch and the SMAD-dependent TGFB/BMP pathway ^26–29^. Our analysis suggests that whereas the presence of RBPJ binding sites in DARs is a sufficient indicator of RO-driven accessibility repression, the presence of SMAD sites is not (Figure 4B left, C). A similar pattern was observed for DEGs associated with DARs containing RBPJ, SMAD, or both binding sites, with genes downregulated after RO treatment predominantly associated with DARs containing an RBPJ binding site (alone or together with SMAD, Figure 4B right, C). These observations suggest that in the endocardium, the downstream effectors of the TGFB/BMP pathway mostly interact with genomic regulatory elements recognized by the Notch effector RBPJ (Figure 4C).

For the positive response to Notch-ligand stimulation, motifs recognized by Activator Protein-1 (AP-1) family members were overrepresented after 6h (Figure 4A, Table S4) ^30^, whereas RUNX and FOX factors were the strongest candidate mediators of the long-term response (Figure 4A, Table S4). RUNX and FOX transcription factors are known to interact with each other^31^ and with Notch^32^.

The three patterns linked to a positive response to Notch activity all show a clear association with C1 DEGs (Figure 3D,E). Accordingly, RBPJ, SMAD, AP-1, and FOX binding sites were the most enriched sites in DARs associated with C1 DEGs (Figure 4F, Table S4). Moreover, consistent with the association of C2 DEGs with regions responding to long term ligand activity (Figure 3D-E), RUNX binding sites were the most enriched sites in DARs associated with C2 DEGs (Figure 4F, Table S4).

Motif enrichment analysis of genomic regions showing increased accessibility after RO inhibition of NICD release suggested that Notch pathway activation represses GATA TF activity in the endocardium (Figure 4A, Table S4). Gata TFs have an active role in the embryonic endocardium^33^. Our observations are consistent with transcriptional repression of Gata factors by the Notch targets Hes and Hey ^34–38^. The analysis of regions showing repressed accessibility after Notch ligand stimulation identified regions recognized by the Hippo pathway member Tead (Figure 4A, Table S4). The Hippo binding site was overrepresented in the negative response to Notch activity in all three conditions, especially Notch ligand stimulation for 24h, with KLF factors being the main candidate drivers of gene expression in this setting (Figure 4A, Table S4). Similar to the interaction between SMAD and RBPJ sites, KLF binding sites were associated with accessibility repression after long-term Notch-ligand stimulation only if they appeared together with TEAD motifs (Figure 4D left, E). The implication of the Hippo pathway in the control of Notch-repressed DEGs is supported by the general downregulation of DEGs associated with DARs containing TEAD sites (with or without KLF motifs in the same genomic element, Figure 4D right, E). Furthermore, TEAD is the most represented binding site, together with GATA, in DARs associated with C3 and C4 DEGs (Figure 4F).

TFs recognizing the overrepresented binding sites showed differential expression, in most cases changing in the same direction as the chromatin changes affecting their potential target genomic elements (Figure 4G), thus indicating an intricate and interdependent regulatory network downstream of NICD. Together, our results propose not only a set of non-coding genomic elements potentially controlling Notch-driven changes in gene expression, but also various cofactors and timeframes for their action as transcriptional regulators.

### Notch-dependent chromatin accessibility changes in the endocardium of embryonic and adult cardiac valves

To assess the importance of the described chromatin accessibility profiles in valve development and homeostasis, we compared our MEEC data with data from E12.5 mouse endocardium^39^ and from human adult mitral valves^17^ (Figure 5A, Table S4). Of the MEEC DARs, 54% overlapped with accessible regions detected in at least one of these in vivo data sets. Interestingly, the percentage of MEEC DARs overlapping with accessible regions increased to 71% when we only considered DARs whose accessibility responded positively to Notch pathway stimulation (clusters a-c, Figure 5B, Table S4), and only 41% of Notch-repressed MEEC DARs were found in vivo (clusters d-g, Figure 5B, Table S4). Based on these observations, we can infer that the role of Notch as a promoter or repressor of genomic elements in MEEC also operates in the endocardium of developing mouse and adult human valves. Furthermore, MEEC DARs whose accessibility was increased by Notch pathway activity (a-c) were statistically associated with Notch-induced DEGs (C1-C2), but only when the DARs were also detected in vivo (Figure 5C). In contrast, MEEC DARs repressed by Notch (d-g) were overrepresented near Notch-repressed DEGs (C3-C4) only when these DARs were not detected in vivo (Figure 5C). Together, these results strongly support a role for endocardial Notch as an activator of non-coding regulatory elements required for valve development and homeostasis and as a repressor of elements able to induce the expression of undesired genes.

**Figure 5.**
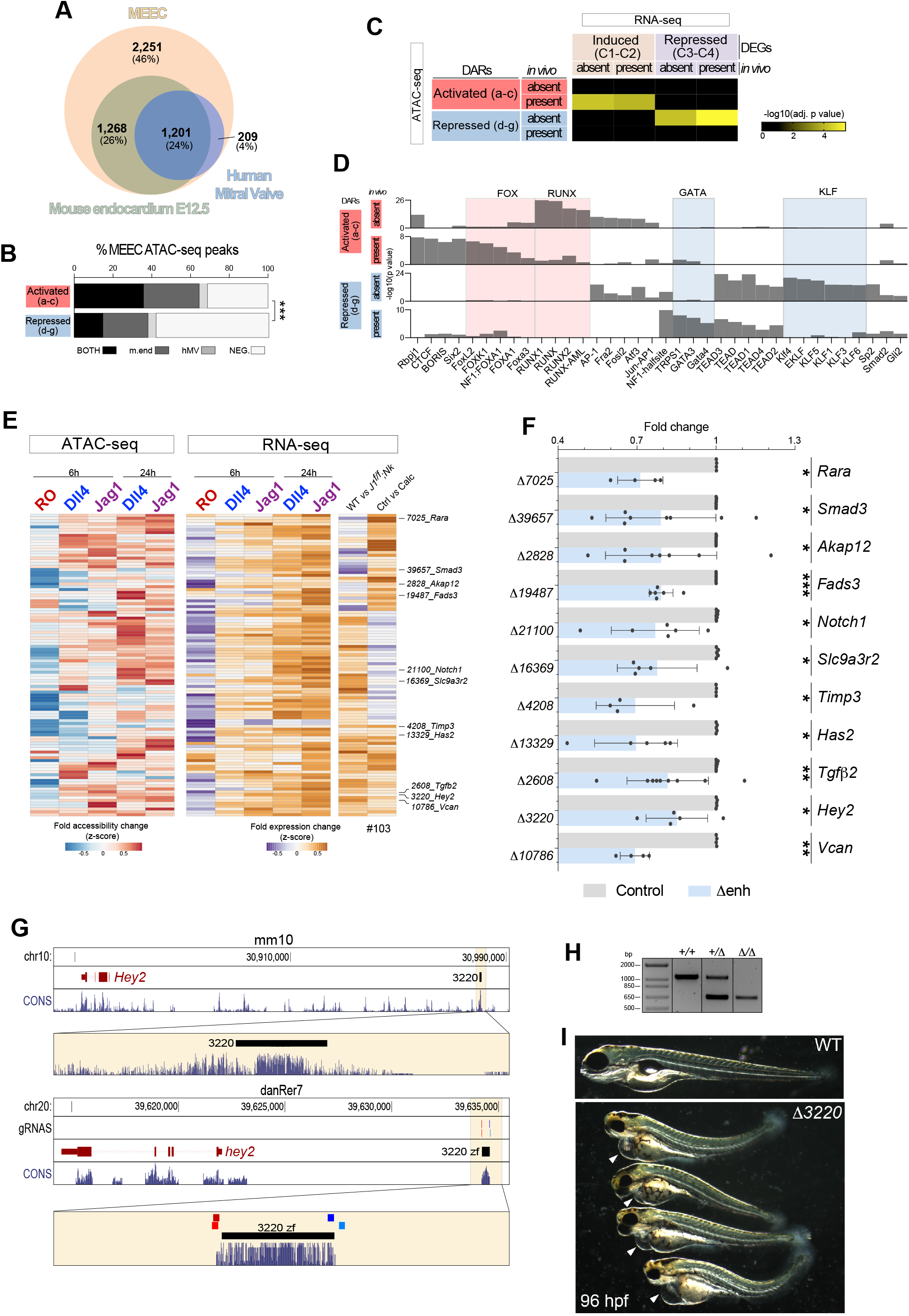
The regulatory potential of Notch-dependent non-coding regulatory elements in mouse embryonic endocardial cells is observed in valve endocardium. (A) Overlap of DARs detected in MEEC with accessible regions identified in the E12.5 mouse endocardium and in adult human mitral valves. (B) Proportion of Notch-activated DARs (clusters a-c, left) and Notch-repressed DARs (clusters d-g, right) overlapping with ATAC-seq peaks detected in human mitral valves, mouse E12.5 endocardium, or both. (C) Enrichment of Notch-activated DARs (clusters a-c) and Notch-repressed DARs (clusters d-g) with proximal DEGs induced (C1-C2) or repressed (C3-C4) by Notch pathway activity. DARs and DEGs are stratified according to overlap (*present*) or non-overlap (*absent*) with peaks detected in vivo. Yellow indicates a significant adjusted p-value. (D) Statistical representation of known binding sites detected in Figure 4A in DARs from clusters a-c and clusters d-g according to overlap (*present*) or non-overlap (*absent*) with peaks detected in vivo. Bars represent the -log10(p value). (E) Heatmap analysis of DAR-DEG pairs where both respond positively to Notch pathway activity and are present in vivo. The heatmaps represent fold accessibility changes in cluster a-c DARs (left), fold expression changes in the associated cluster C1-C2 DEGs (center), and fold expression changes in genes significantly enriched in E14.5 WT versus *Jag1^flox^;Nkx2.5-Cre* semilunar valves and control versus calcified adult human valves (right). Gene names to the right identify candidate endocardial Notch target genes putatively regulated via the associated non-coding genomic elements (DARs). (F) qRT-PCR expression analysis of the candidate target genes in control MEEC (transfected with the empty vector) and MEEC transfected with the vector containing the guide RNAs specific for selected DARs. Deleted enhancers are indicated on the left, genes assayed on the right. * p<0.05, ** p<0.01, *** p<0.001 (paired t-test). (G) Mouse and zebrafish genome maps indicating the location of the Notch-dependent regulatory element (*3220*) associated with *Hey2* and location of the guide RNAs used to delete this element in zebrafish. (H) PCR of genomic DNA from three 96 hpf zebrafish siblings (wild type, element Δ*3220* mutant heterozygote, and Δ*3220* null mutant). (I) Bright-field images of a sibling control and 4 Δ*3220* null mutant embryos at 96 hpf. Arrowheads indicate the pericardial edema.

The integration of in vitro and in vivo data also helped us to identify cofactors with a potential role in valve endocardium. For example, in contrast to FOX, RUNX binding sites were specifically represented in those MEEC DARs activated by Notch and undetected in mouse developing endocardium or adult human valves (Figure 5D), suggesting that coordinated activity of NOTCH and RUNX in the endocardium is not involved in valve development. For DARs repressed by Notch, KLF motifs were specifically overrepresented if the DARs were also repressed in valves (Figure 5D), suggesting that repression of KLF TFs, together with TEAD, in the endocardium may be necessary for valve development and homeostasis.

Based on this analysis, we generated a list of high-confidence non-coding genomic elements that are detected in vivo, respond positively to Notch-pathway activation in the embryonic endocardium, and show an association with endocardial Notch target genes in the in vivo data (Figure 5E, Table S5). Based on their biological interest, selected elements were validated for their ability to regulate the expression of their putative target genes (Figure S4A). Using CRISPR-Cas9 gene editing technology, we transfected MEEC with a pair of specific guide RNAs targeting each genomic element (Figure S4A, Table S5). Although deletion of the genomic element occurred in only a fraction of the transfected cells (Figure S4B), this resulted in a significant reduction in the expression of the putative target genes (Figure 5F). This was the case for genomic elements associated with genes downregulated in calcified human valves (*Rara*, *Smad3*, *Akap12*, *Fads3*), in *Jag1^flox^;Nkx2.5-Cre* E14.5 semilunar valves (*Notch1*, *Slc9a3r2*), and in both models (*Timp3*, *Has2*, *Tgfb2*, *Hey2*, *Vcan*). In this last group, the *Hey2*-associated genomic element *3220* showed high sequence conservation as well as conserved proximity to the putative target *Hey2* (Figure 5G). We used 2 pairs of guide RNAs (Figure 5G, Table S5) to generate a knock-out zebrafish model for the *3220* element (Figure 5H). The progeny of the Δ3220 line showed pericardial distension, an unlooped heart, and a bent tail at 96 hours post fertilization (hpf), a similar phenotype to that of *hey2* loss-of-function models ^40–42^ (Figure 5G). Our in vitro and in vivo validation experiments thus confirm the functional relevance of the identified regions.

### Notch and Hippo signaling pathways converge in the endocardium during heart valve development

The integration of data on transcriptomic and epigenomic changes in MEEC after Notch manipulation revealed a strong association between inhibition of gene expression and repression of genomic elements potentially recognized by TFs of the TEAD family. Based on this finding, we sought evidence for cooperation between the Notch and Hippo pathways in the embryonic endocardium. Genes targeted by the Hippo pathway downstream effectors Yes-associated protein (YAP) and transcriptional co-activator with PDZ-binding motif (TAZ) were identified in available transcriptional data from human umbilical endothelial cells (HUVEC) transduced with the transcriptionally active forms YAP^S127A^ and TAZ^S89A^ ^43^. We then assessed the representation of these YAP/TAZ endothelial target genes among the MEEC DEGs identified in the MEEC–OP9 co-culture model (clusters C1-C4). DEGs repressed by Notch signaling activation (C3-C4 DEGs) were statistically enriched in YAP/TAZ-positive target genes (Figure 6A), suggesting that NOTCH represses YAP/TAZ transcriptional activity. This antagonism between endocardial NOTCH and YAP/TAZ was also observed in the mouse in vivo data, where genes repressed by Notch activation in E14.5 semilunar valves were enriched in YAP/TAZ-positive target genes (Figure 6B). Analysis to determine if this regulation could be mediated by genomic elements repressed by Notch activity (clusters d-g, Figure S3A) showed that Notch-repressed YAP/TAZ-positive target genes in the endocardium were mostly associated with genomic elements repressed by Notch activity not only in vitro but also in vivo (Figure 6C). In addition, the presence of the specific binding motif for the YAP/TAZ interactor TEAD was strongly predicted by Notch-repressed accessibility in vitro and in vivo and association with genes repressed by Notch and induced by YAP/TAZ (Figure 6D). Together, this evidence suggests that endocardial Notch acts as a repressor of YAP/TAZ-dependent non-coding elements, resulting in the subsequent inhibition of direct target genes. We considered that this repression might be accompanied by a reduction in the content of Yap1 in the nucleus. Analysis of the subcellular localization of Yap1 in MEEC revealed that exposure to recombinant JAG1 significantly reduced nuclear Yap1 content, whereas exposure to RO increased it (Figure 6E-F). Like Notch, endocardial YAP1 is required for myocardial proliferation ^44^ and cellularization of the valve endocardial cushions ^45^. Our results reveal convergence of Hippo and Notch signaling and suggest that Notch acts as a modulator of Yap1 activity in the embryonic endocardium (Figure 6G).

**Figure 6.**
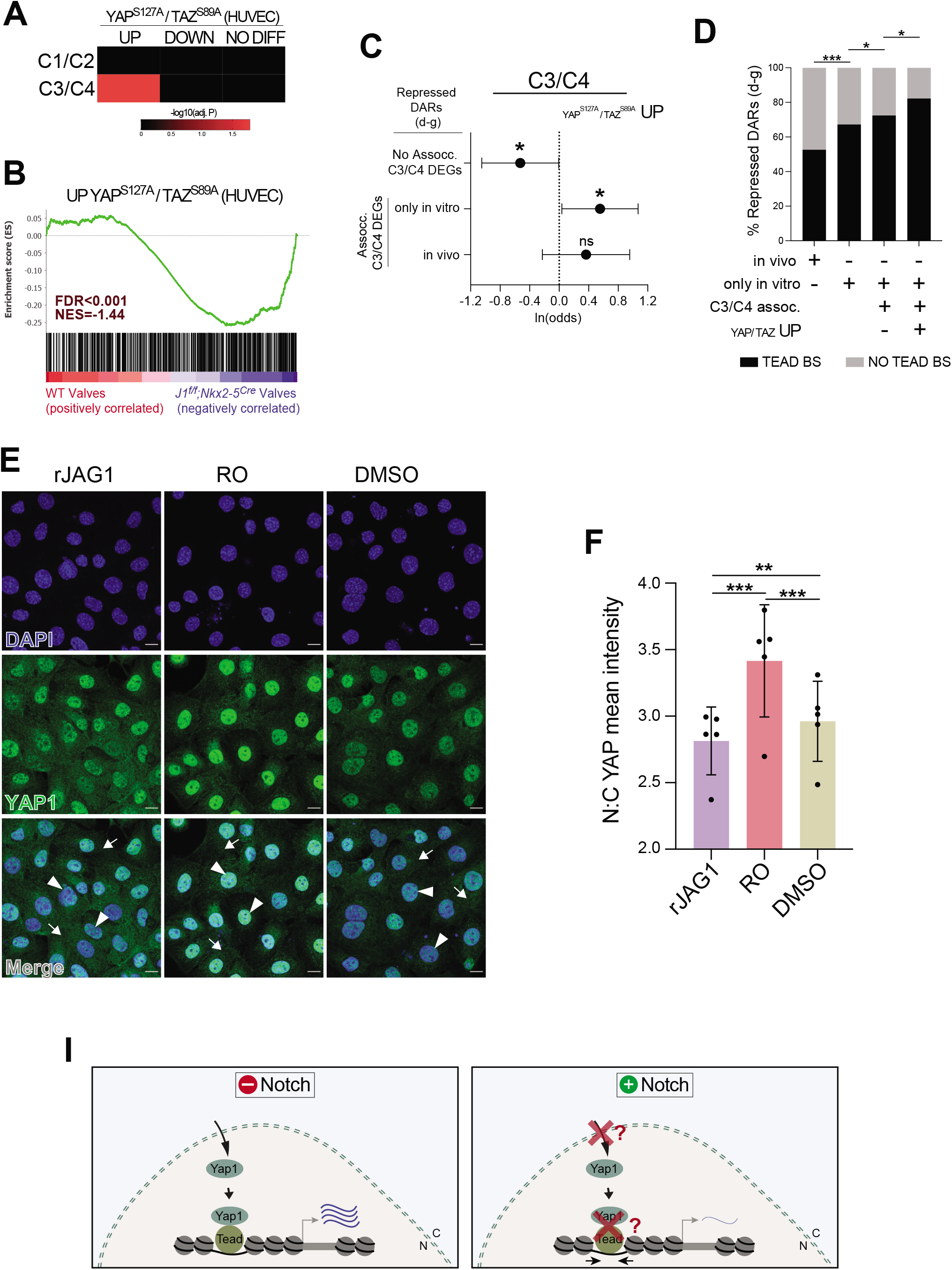
The Hippo and Notch signaling pathways converge in mouse embryonic endocardial cells. (A) Enrichment of genes upregulated, downregulated or with no change in HUVEC after YAP^S127A^/TAZ^S89A^ transfection, in DEGs induced (C1-C2) or repressed (C3-C4) by Notch-pathway activity. Red indicates a significant adjusted p value. (B) Gene-set enrichment analysis showing the distribution of YAP^S127A^/TAZ^S89A^-positive target genes in HUVEC within the ranking of all genes identified in E14.5 WT and *Jag1^flox^;Nkx2.5-Cre* semilunar valves. FDR values represent the statistical significance of the enrichment score. NES, normalized enrichment score. (C) Odds ratios for finding YAP/TAZ-positive target genes in C3-C4 DEGs showing no association or an association with Notch-repressed DARs, either with or without overlap with accessible genomic regions detected in developing mouse or adult human valves. * p<0.05, ** p<0.01, *** p<0.001 (Fisher’s exact test). (D) In Notch-repressed DARs, proportion of DARs containing TEAD binding sites (BS) under different conditions, such as overlap or no overlap with accessible genomic regions detected in the developing mouse endocardium or adult human valves, association with C3-C4 DEGs, or association with C3-C4 DEGs induced by YAP/TAZ activity. * p<0.05, ** p<0.01, *** p<0.001 (Fisher’s exact test). (E) Immunostaining of YAP1 in MEEC exposed to DMSO, RO, or recombinant JAG1. Nuclei are counterstained with DAPI. (F) Quantification of the nuclear:cytoplasmic intensity ratio of YAP1 in MEEC exposed to DMSO, RO, and rJAG1. * p<0.05, ** p<0.01, *** p<0.001 (two-way ANOVA with multiple comparisons by Tukey’s correction). (G) Proposed interaction between Notch and Yap/Taz in the endocardium.

## Discussion

In this study, we stimulated MEEC with Jag1 and Dll4 ligands expressed on OP9 cells to identify the transcriptomic and epigenomic consequences of NOTCH manipulation. Our analysis defines a primary (short-term) and a secondary (long-term) transcriptional response to NOTCH signaling activation and identifies a set of candidate transcription factors that potentially act in a well delimited time window. We confirmed the relevance of these findings through two approaches: 1) incorporation into our analysis of in vivo transcriptomic and epigenomic data from developing valves of wild type and Notch mutant mice and from control and diseased adult human heart valves; 2) validation of the regulatory potential of a selection of the identified genomic elements against the in vitro and in vivo data.

During cardiac valve development, the Notch1 receptor is sequentially activated in the endocardium by the ligands Dll4 and Jag1. At early stages, Dll4 and Jag1 are expressed in valve endocardium, but only Dll4 is required for the initiation of valve formation through EMT. Once the valve primordia are well formed, valve morphogenesis requires Jag1 expressed in the myocardium ^6^, and a more recent study found that Jag2 is also required ^46^. Interestingly, during early valvulogenesis, manic Fringe (MFng), a member of the Fringe family of glycosyltransferases that elongate carbohydrates attached to the Notch extracellular EGF-like repeats, is expressed in the endocardium, and its expression is progressively reduced as development proceeds ^6, 20^. In Drosophila, Fng favors Delta-Notch signaling and attenuates Jag/Ser–Notch signaling ^47, 48^. In a test of this Fng selectivity in MEEC, luciferase-based assays and qPCR determination of the transcription of a bona fide Notch target gene revealed markedly enhanced Dll4 signaling in the presence of MFng ^20^. Our comparison of the transcriptional output after Dll4 and Jag1 signaling to MEEC, especially after short-term stimulus, showed Dll4 to be less efficient than Jag1 at inducing a transcriptional response in target cells. Our transcriptional analyses in MEEC did not detect *MFng* expression, strongly suggesting that the differences in transcriptional output triggered by the two ligands simply reflect this lack of Fng and that there is no NOTCH ligand-specific transcriptional response in the endocardium.

The nuclear translocation of NICD after NOTCH ligand–receptor binding allows its interaction with the transcriptional repressor RBPJ, releasing corepressors and recruiting the coactivator Mastermind (MAML) to give rise to a transcriptional activator complex that initiates the Notch-dependent transcriptional response. Besides the well-known NOTCH target genes, which include the basic–helix–loop–helix transcriptional factors of the Hes and Hey families ^27, 49, 50^, several other genes have been proposed to be Notch-dependent in the endocardium during cardiogenesis ^6, 20, 27, 51–55^. However, it remains unclear what position these genes occupy in the Notch-dependent transcriptional landscape and what cofactors are required for their endocardial expression. Our combined use of the γ-secretase inhibitor RO in unstimulated MEEC and two time windows of Notch-ligand stimulation revealed the existence of a primary and sustained “early” transcriptional response and a secondary “late” response in the embryonic endocardium. The primary response is triggered by the baseline activity of Notch ligands and gradually increases with ligand stimulation for 6 and 24 hours. Together with RBPJ, SMAD binding sites were the most represented motifs in genomic regions repressed by RO. Smad activity in the endocardium is required to initiate EMT and is induced by TGFβ2 and BMP2 signaling in the AVC and OFT territories^27, 29, 51, 56, 57^. *Tgf*β*2* is expressed in the presumptive valve endocardium and myocardium and is NOTCH-responsive in vivo ^27, 51^. In the MEEC–OP9 co-culture assay, where there is no signaling from the myocardium, *Smad3* and *Tgf*β*2* expression (and TGFβ2 secretion) was induced by Notch-pathway activation through the activity of at least one specific genomic regulatory element in each case. The presence of RBPJ binding sites in most potential Smad-dependent regulatory elements suggests a crucial role for Notch not only as a TGFβ2 activator, but also as a required cofactor in the transcriptional induction of target genes. Further studies will be needed to determine if myocardial TGFβ/BMP signaling to the endocardium operates through Notch-independent genomic elements.

Genomic regions with increased chromatin accessibility after short-term stimulation were enriched in binding sites recognized by Activator protein 1 (AP-1), a transcription factor heterodimer composed of proteins from the c-Fos, c-Jun, ATF, and JDP families. AP-1 proteins have been proposed to act in parallel with Notch in neurons and in cancer ^30, 58, 59^, with AP-1 playing an active role in the endothelium ^60, 61^. The early positive response to Notch-pathway manipulation of *Jun* and *Fosb* (both in cluster C1) suggests these proteins as Notch partners in the transcriptional response of the embryonic endocardium. Similarly, *Foxc1*, expressed in the endocardium and required for outflow-tract morphogenesis ^62^, may recognize the abundant FOX binding sites in genomic regions induced by long-term Notch-ligand stimulation. Overall, NOTCH-responsive genomic elements containing these binding sites showed a significant association with C1 DEGs, indicating a sequential activity of specific transcription factors acting cooperatively in response to Notch-pathway activation.

Mutations in NOTCH1 have been linked to an increased risk of BAV and aortic valve calcification ^10, 63–66^ . This strongly justifies our choice of in vivo datasets to give biological context to the Notch-dependent gene expression changes detected in MEEC: a mouse model of disrupted cardiac NOTCH signaling that develops BAV; and adult valves from healthy controls and CAVD patients. Both datasets showed evidence of the primary and secondary positive transcriptional responses to Notch signaling (expression clusters C1 and C2), with the primary response (C1) more prominently impaired in *Jag1^flox/flox^;Nkx2-5-Cre* valves and the secondary response (C2) more downregulated in human CAVD. The good representation of Notch-responsive endocardial genes in *Jag1^flox/flox^;Nkx2-5-Cre* semilunar valves, together with the associated genomic elements found in mouse and human valves, allowed us to propose a set of genes and genomic elements potentially involved in semilunar valve development. In contrast to the mouse model, human samples were not selected based on Notch dysregulation. Nevertheless, many of the genes responding positively to Notch (especially long-term stimulation) were downregulated in CAVD samples. Putative RUNX-dependent genomic elements were overrepresented near genes with a long-term positive response to Notch activity (C2), although these elements were mostly inaccessible in mouse and human valves. *Runx2* derepression has been suggested to induce CAVD in NOTCH1 haploinsufficient mice^10^, but our MEEC-OP9 co-culture system confirmed Notch as an inducer of *Runx2* expression in the embryonic endocardium, similar to its action in chondrocytes ^67^. In line with our observation, a more complex pro-osteogenic and inflammatory program was proposed to be repressed by Notch endothelial activity ^23^. In the MEEC–OP9 assay, *Runx1*, another RUNX family member, showed a positive response to Notch activation. RUNX1 is also involved in bone formation^68, 69^, plays crucial roles in the hemogenic endothelium^70^ and in the endocardium^71^, and has been described as a Notch target gene in hematopoiesis^72^ and T cell development^73^. The Notch-dependent transcriptomic signature in CAVD proposed here is associated with metabolic activity and transcriptional regulation, which could be related to a slowed down metabolic status in calcified valves. Rather than the proposed role of NOTCH as a repressor of calcification, we suggest that by converging with other pathways, Notch regulates a more complex program to maintain valve homeostasis.

In addition to the positive transcriptional response, Notch activity also repressed a set of distal genomic elements in the MEEC-OP9 co-culture system that were also mostly repressed in the developing mouse endocardium and mitral human valves, consistent with the proposed role of Notch in valve development and homeostasis ^74^. A common feature of these elements is the presence of TEAD binding sites, indicating an association with genes downregulated by Notch activity. Tead transcription factors require the activity of co-factors such as the Hippo signaling pathway effectors Yap and Taz ^75^. The Hippo signaling pathway is implicated in the control of organ growth ^76^, operating through the serine/threonine kinase Mst1/2 and the downstream kinase Lats1/2, which phosphorylates the transcription co-factors Yap and Taz ^77^. When unphosphorylated, Yap and Taz translocate to the nucleus, where they recognize the cofactor Tead. Cardiac Yap/Taz activity has been implicated in the regulation of cardiomyocyte proliferation ^78^. In the endocardium, the relationship between Yap1 and Notch is unclear. Endocardial ablation of both Yap1 and Notch activity results in a hypoplastic myocardium, due at least in part to the downregulation of the growth and differentiation regulator Neuregulin 1 ^44, 52^. In the presumptive valve territory, genetic ablation of *Yap1* or *Notch1/Rbpj* impairs valve formation, resulting in hypocellular endocardial cushions and the downregulation of the EMT driver *Snai1* ^45, 51^. However, whereas endothelial deletion of *Yap1* reduces endocardial proliferation ^45^, Notch signaling ablation results in a hyperproliferative endocardium ^52^, suggesting that the YAP1 chromatin targets that promote endocardial proliferation may become active after NOTCH inactivation. Our results identify Notch as a negative modulator of YAP/TAZ activity. In addition to repressing genomic elements potentially recognized by YAP/TAZ, Notch-pathway activation inhibits many endothelial targets of YAP/TAZ activity in the embryonic endothelium, both in vitro and in vivo. In vitro repression of endocardial Notch nuclear translocation resulted in an increase in nuclear Yap1, suggesting that the Notch and Hippo pathways may converge in the endocardium at the level of Yap1 nuclear translocation. Future in-depth analysis will be needed to uncover the mechanisms underlying the functional relationship between NOTCH and YAP in the embryonic endocardium and its relevance to cardiac development and disease.

## Supporting information

Supplemental file containing Suppl. methods, Suppl. Figures and Legends & Suppl. Tables legends.

## Acknowledgments

We thank E. Diaz for zebrafish husbandry; the CNIC Flow Cytometry Unit for FACS; the CNIC Genomics Unit for RNA-seq; CNAG-CRG for high throughput sequencing for ATAC-seq; the CNIC Bioinformatics Unit for RNA-seq and ATAC-seq data analysis; P.A. Morocho, S. Urra, and D. Muñoz for experimental work; and S. Bartlett for English editing.

## Nonstandard Abbreviations and Acronyms

ATAC-Seq: assay for transposase-accessible chromatin with sequencing
AVC: atrioventricular canal
BAV: bicuspid aortic valve
CAVD: calcific aortic valve disease
DAR: differentially accessible region
DEG: differentially expressed gene
ECM: extracellular matrix
EMT: epithelial-to-mesenchyme transition
GO: gene ontology
HUVEC: human umbilical endothelial cells
MEEC: mouse embryonic endocardial cells
OFT: outflow tract
RO: RO4929097, a γ-secretase inhibitor
TAV: tricuspid aortic valve
YAP: yes-associated transcriptional protein
TAZ: transcriptional co-activator with PDZ-binding motif

