## Supplemental file containing Suppl. methods, Suppl. Figures and Legends & Suppl. Tables legends. for "A cooperative response to endocardial NOTCH signaling stimulation regulates transcriptional activity during cardiac valve development and disease"

**ONLINE SUPPLEMENTARY DATA**

---

**Detailed methods**

### Cell Culture

MEEC were cultured on 0.1% gelatin-coated dishes and maintained in Dulbecco's modified Eagle's medium (DMEM) (Gibco) supplemented with 10% fetal bovine serum (FBS) and 1% penicillin/streptomycin. GFP-, Jag1- or Dll4-expressing OP9 cells were cultured on 0.1% gelatin-coated dishes and maintained in MEM Alpha (1X) (Gibco) supplemented with 20% fetal bovine serum (FBS) and 1% penicillin/streptomycin. Cultures were incubated at 37°C in a humidified atmosphere containing 5% CO2.

**MEEC-OP9 coculture**

To define the effect of Dll4- and Jag1-mediated cell-to-cell signaling, MEEC were co-cultured for 6h (50% confluency) or 24h (25% confluency) over GFP-, Jag1- or Dll4-expressing OP9 cells at 90% confluence in DMEM supplemented with 10% fetal bovine serum (FBS) and 1% penicillin/streptomycin. To define the effect of inhibiting NICD nuclear translocation, MEEC were cultured with DMSO (vehicle) or 20 µM of the γ-secretase inhibitor RO4929097 for 24 h, and the MEEC were then co-cultured for 6h (50% confluency) over GFP-expressing OP9 cells (90% confluency) in DMEM supplemented with 10% fetal bovine serum (FBS), 1% penicillin/streptomycin, and renovated DMSO or RO4929097. All cells were then trypsinized, washed in PBS, and harvested for FACS sorting (GFP-negative selection).

**MEEC culture with recombinant ligands**

For RNA-seq, the recombinant NOTCH ligands DLL4-His (R&D Systems 1389-D4-050) and JAG1-Fc (R&D Systems 599-JG-100) were attached to 24-well dishes as described [1]. Once MEEC were seeded, DMSO was added to control and recombinant-ligand stimulated cells and 20 µM RO4929097 (Selleck Chemicals S1575) was added to designated wells to inhibit Notch activity. After 24 h at 37°C, cells were washed twice with PBS and incubated for an additional 14 h in serum-free DMEM together with RO4929097 or DMSO as appropriate. Each condition was performed in triplicate.

**CRISPR editing**

The sgRNA pairs for the desired deletions were designed in (crispor.tefor.net) (Table S5). Each sgRNA pair was cloned in pDG458 (Addgene #100900) according to the provided *pDG* One-Step Cloning Protocol* (www.addgene.org/100900). After transformation and bacterial selection with ampicillin, DNA was recovered with the Nucleospin Tissue Kit (Macherey-Nagel #740952.50).

**Cell culture and transfection**

MEEC at 70-80% confluence were transfected with 15 ug of pDG458 vector, either empty or carrying the sgRNA pair, using Lipofectamine LTX Reagent with PLUS Reagent (ThermoFisher #15338100). A mix containing the plasmid and the LTX and PLUS reagents was added to the medium, and cells were incubated for 48 hours with a change of medium after 24 hours. Cells were then sorted (GFP+) and frozen.

**Gene expression analysis**

Total RNA from cells was extracted with the PicoPure RNA Isolation Kit (ThermoFisher #KIT0204). cDNA was synthesized using the High-Capacity cDNA Reverse Transcription Kit with RNase Inhibitor (ThermoFisher Scientific #4374966). qPCR was performed with the PowerSYBR green PCR Master Mix (Applied Biosystems #4367659). Primers (Table S5) were designed using the NCBI Primer design tool (ncbi.nlm.nih.gov/tools/primer-blast). The Delta-Delta Ct Method was used to analyze gene expression differences between recombinant JAG1-stimulated MEEC transfected with pDG458-sgRNA or pDG458. A paired t-test was performed using the $\Delta$Ct values in Prism 9.

**RNA-seq**

Total RNA from cells was extracted with the PicoPure RNA Isolation Kit (ThermoFisher #KIT0204) and was used to generate barcoded RNA-seq libraries with the NEBNext Ultra RNA Library preparation kit (New England Biolabs). Briefly, poly A+ RNA was purified using poly-T oligo-attached magnetic beads followed by fragmentation and then first and second cDNA strand synthesis. Next, cDNA 3' ends were adenylated and the adapters were ligated followed by PCR library amplification. Finally, the size of the libraries was checked using the Agilent 2100 Bioanalyzer DNA 1000 chip, and their concentration was determined using the Qubit® fluorometer (Life Technologies). Sequencing reads were pre-processed by means of a pipeline that used FastQC (bioinformatics.babraham.ac.uk/projects/fastqc) to assess read quality and Cutadapt [2] to trim sequencing reads (eliminating Illumina adaptor remains) and to discard reads shorter than 30 bp. Resulting reads were mapped against reference transcriptome GRCm38.91 and quantified using RSEM [3]. Percentages of reads participating in at least one reported alignment were around 82%. Expected expression counts calculated with RSEM were then processed with an analysis pipeline that used the Bioconductor package Limma [4] for normalization (using TMM method) and differential expression testing, taking into account only those genes expressed with at least 1 count per million (CPM) in at least three samples and , when required, using a random variable to define pairs of samples with the same origin. Changes in gene expression were considered significant if associated with a Benjamini and Hochberg adjusted p-value $< 0.05$.

Mouse ensemble IDs were converted to human ensemble IDs using biomart (www.ensembl.org). Heatmaps were generated with ClustVis (biit.cs.ut.ee/clustvis_large) [5].

Gene ontology enrichment analysis of biological processes terms was performed with Metascape (metascape.org) [6]. As background, we used the genes detected by RNA-seq in MEEC cocultured with OP9 cells.

Gene set enrichment analyses were performed with GSEA 4.2.1 [7] [8] on the complete set of expressed genes in WT vs *Jag1$^{flox/flox}$; Nkx2-5-Cre* mouse semilunar valves [9], adult human control and calcified valves [10] vs C1-C4 DEGs (in Figure 2), and WT vs *Jag1$^{flox/flox}$; Nkx2-5-Cre* valves against differentially expressed genes in HUVEC transduced with $YAP^{S127A}$ and $TAZ^{S89A}$ [11].

**Interactome prediction**

Genes in clusters A1-A3 served as input node data for the STRING 11.5 database (string-db.org/)[12]. To obtain more stringent interactions, the *Textmining* option in STRING was excluded, and the minimum required interaction score considered was 0.7 (high confidence). Resulting networks were then replotted using Cytoscape 3.9.1 [13] applying the Attribute Circle Layout setting for each cluster individually.

**ATAC-seq**

Three independent ATAC-seq experiments were performed. 1) MEEC cultured for 6 hours over control OP9 cells or OP9 cells expressing Dll4 or Jag1; 2) MEEC cultured for 24 hours over control OP9 cells or OP9 cells expressing Dll4 or Jag1; 3) MEEC cultured for 6 hours over control OP9 in the presence of DMSO (vehicle) or RO4929097. Each experiment was replicated 4 times.

ATAC-seq was performed as described in [14]

Bioinformatics analysis: For each experiment, sequencing reads were pre-processed by means of a pipeline that used FastQC (bioinformatics.babraham.ac.uk/projects/fastqc) to assess read quality and Cutadapt [2] to trim sequencing reads (eliminating Illumina and Nextera transposase adapter contaminations) and to discard reads shorter than 30 bp. Resulting reads were then mapped against reference genome GRCm38/MM10 with a pipeline that used bowtie2 [15] as aligner; Piccard (broadinstitute.github.io/picard) to mark duplicate alignments; and samtools [16] to eliminate duplicate, chimeric, and sub-optimally multi-mapped alignments, thus keeping only properly paired and mapped reads. Alignments against the mitochondrial genome and chromosome Y were also removed. Once filtered alignments were obtained, peaks (accessible DNA regions) were called with MACS2 [17], using parameters "--nomodel --shift -100 --extsize 200", and "-q 0.05" as the false discovery rate cut-off. Next, filtered alignments and peaks (in bam and xls formats, respectively) were processed with the R package DiffBind to define a consensus peak set, recalculate and normalize peak coverage across samples, and identify differentially accessible regions (DARs), using EdgeR as the analysis method. Due to the low number of DARs identified with FDR < 0.05, a very relaxed analysis was performed. For each independent experiment, fold accessibility changes were z-scored, and DARs were defined as regions deviating more than 2 standard deviations and with a raw p-value < 0.0455 (corresponding to a z-score > 2 or < -2).

A final consensus interexperiment set of peaks was defined with the R package DiffBind, considering as DARs the consensus peaks overlapping with the DARs defined in each individual experiment. Heatmaps were generated with NG-CHM BUILDER (build.ngchm.net) [18].

The association of differential accessibility ATAC-seq peaks with proximal differential expressed genes was analyzed with GREAT (version 4.0.4) (www.great.stanford.edu) [19], using the GRCm38 assembly and the default “Basal plus extension” (Figure 3, 4D, 5C) and “Single nearest gene” settings (Figure 4E, 5B).

ATAC-seq coordinates were converted to different assemblies using the liftOver tool on the UCSF Genome Browser [20], with minimal ratios of remapped bases of 0.95 for mouse-mouse conversions and 0.5 for mouse-human conversions.

Intersection and merging of genomic coordinates was performed with the Table Browser tool on the UCSC Genome Browser and the Galaxy platform [21] on the Mississippi server (mississippi.sorbonne-universite.fr).

Enrichment analysis for differential accessibility peaks was performed with GSEA 4.2.1 on the complete set of consensus ATAC-seq peaks against differential accessibility ATAC-seq peaks associated with C1-C4 DEGs clusters.

**Correlation of expression clusters and chromatin accessibility patterns**

Chromatin accessibility pattern—expression pattern (Figure 5C) and expression pattern—expression pattern (Figure 6A) significant enrichments were identified by computing a standardized Pearson residual for every pattern. This corresponds to a $\chi 2$ test. The p-value associated with each standardized Pearson residual was calculated, and a Bonferroni correction was performed to adjust for multiple tests.

**Transcription factor analysis**

Known motif overrepresentation in differential accessibility ATAC-seq peaks from each group was analyzed with the findMotifsGenome.pl script in the HOMER (v3.12) collection [22], using the complete set of consensus ATAC-seq peaks as background. The genomic locations of differential accessibility ATAC-seq peaks were annotated using annotatePeaks.pl script.

**Zebrafish husbandry and transgenic lines**

Zebrafish were raised under standard conditions at 28$^{o}$C [23]. We used the *Tg(fli1a:GFP)* transgenic line [24].

**CRISPR/Cas9 deletion of the zebrafish *hey2* cis-regulatory element**

The *hey2* cis-regulatory element (CRE) of was deleted by targeting two different guide RNAs bilateral to each region. The guide RNAs were generated as described [25],[26]. Briefly, the oligos (Table S5) were inserted into the pX330 vector [27], which had been linearized with BbsI (New England Biolabs, Ipswich, MA). The guide RNAs were amplified with forward primers containing the T7 polymerase promoter-specific sequences (Table S5) and the reverse primer AAAAAGCACCGACTCGGTGCCA. The guide RNAs were injected into one-cell stage *Tg(fli1a:GFP)* embryos [24] together with the Cas9 protein (NEB). Animals carrying the CRE deletion were identified by PCR using the primers listed in Table S5. The PCR product was confirmed by an agarose gel electrophoresis, and the mutant animals were identified by the size difference of the PCR products.

**Tissue processing**

Dissected E14.5 embryos were fixed in 4% PFA O/N for *in situ* hybridization (ISH). After dehydration through a graded ethanol series followed by xylene washes, embryos were embedded in paraffin at 65ºC.

**In situ hybridization**

ISH was performed as described [28]. Details of probes will be provided on request.

**Immunostaining**

For immunocytochemistry, MEEC were seeded on 12 mm glass coverslips (VWR 631-1577P) in 24-well plates at a density of 70,000 cells per well. The conditions for Notch stimulation with recombinant JAG1 and Notch inhibition were as described above. After stimulation for 24h, cells were fixed in 4% PFA for 15 min at RT, washed 3 times with PBS, permeabilized with 0.2% Triton X-100 in 1% bovine serum albumin (BSA) for 5 minutes, and blocked with FBS during 1h. Cells were incubated overnight at 4°C with anti-YAP1 (1:100; Santa Cruz Biotechnology #sc-101199), followed by incubation with goat anti-mouse Alexa Fluor 568 (1:250; Thermofisher #A11031).

**Confocal imaging**

Confocal images of stained cells and embryo sections were acquired with a Leica SP5 laser-scanning confocal microscope and with a Leica gated STED-3X-WLL SP8 laser-scanning confocal microscope, respectively. Alexa568 signals were obtained with a 566 nm laser, fluorescein signals with a white nm laser, and DAPI with a 405 nm diode laser. Images were collected as Z-stacks with 1 µm spacing between each plane. Section thickness was 2.014 µm for cells and 8 µm for embryos. Images were processed with Fiji ImageJ 1.32J.

To quantify YAP1 subcellular localization, we used an ImageJ macro to calculate the nuclear-to-cytoplasmic ratio (N:C) from the mean YAP1 fluorescence intensities in the nucleus and cytoplasm after subtracting the fluorescence background. First, a maximum intensity projection of the 3-plane images was obtained, followed by the subtraction of the background signal in the red channel using Huang's algorithm as a threshold. To avoid bias, nuclear and cytoplasmic regions of each cell were delimited through the DAPI channel. Cytoplasmic regions were defined as the perinuclear region extending 2 µm from the nucleus boundary. Cells whose boundaries overlapped were excluded from the analysis. To derive N:C ratio, the signal in the nuclear ROI was divided by the signal in the perinuclear ROI. Differences were considered statistically significant at a p-value $<0.05$ (two-way ANOVA, multiple comparisons with Tukey's correction).

**Data representation**

Prism 9 and BioVenn (biovenn.nl) were used for charts, heatmaps (except for RNA-seq and ATAC-seq), and Venn diagrams.

**Accession numbers**

Data are deposited in the NCBI GEO database under accession number GSE223735.

RNA-seq from E14.5 WT and *Jag1$^{flox/flox}$;Nkx2-5-Cre* micro-dissected semilunar valves: GSE74556. RNA-seq from adult control and calcified bicuspid (BAV) and tricuspid aortic valves (TAV): GSE148219. ATAC-seq from E12.5 mouse endocardial lineage: E-MTAB-3972. ATAC-seq from human mitral valves: PRJNA690001

### Supplementary Figures and Legends

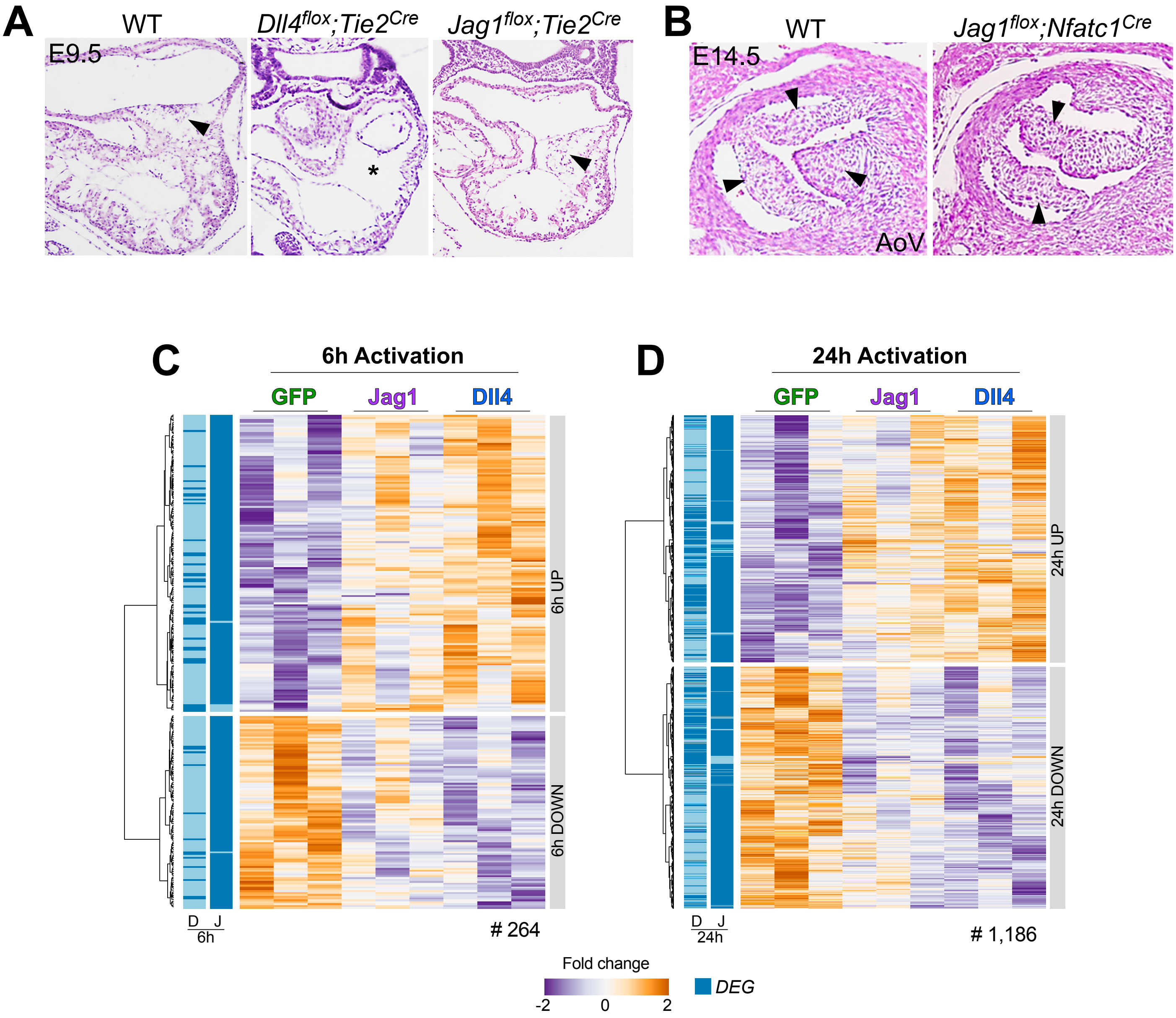


**E**

| | Dll4 6h | Jag1 6h | Dll4 24h | Jag1 24h |
|---|---|---|---|---|
| Dll4 6h | 1.000 | 0.784 | 0.610 | 0.463 |
| Jag1 6h | 0.784 | 1.000 | 0.451 | 0.572 |
| Dll4 24h | 0.610 | 0.451 | 1.000 | 0.691 |
| Jag1 24h | 0.463 | 0.572 | 0.691 | 1.000 |

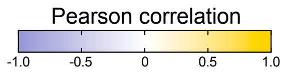


Luna-Zurita et al._Figure S1

**Figure S1. Transcriptional effects of Dll4 and Jag1 activity in the embryonic endocardium**

(A) Transverse hematoxylin and eosin (H&E)–stained sections from E9.5 WT, *Dll4*$^{flox}$*;Tie2-Cre*, and *Jag1*$^{flox}$*;Tie2-Cre* hearts. Arrowheads indicate invading mesenchymal cells, and the asterisk marks the lack of invading mesenchymal cells in the atrioventricular canal.

(B) Transverse H&E–stained sections from E14.5 WT and *Jag1*$^{flox}$*; Nfatc1-Cre* aortic valves. Arrowheads indicate the three leaflets in WT aortic valves and the two leaflets in *Jag1*$^{flox}$*; Nfatc1-Cre* aortic valves.

(C) Hierarchical clustering of genes showing differential expression in mouse embryonic endocardial cells co-cultured for 6 hours with OP9-DLL4 and OP9-JAG1 cells.

(D) Hierarchical clustering of genes showing differential expression in mouse embryonic endocardial cells co-cultured for 24 hours with OP9-DLL4 and OP9-JAG1 cells.

(E) Pearson correlation between the changes in MEEC gene expression induced by co-culture with OP9-DLL4 and OP9-JAG1 cells for 6 and 24 hours.

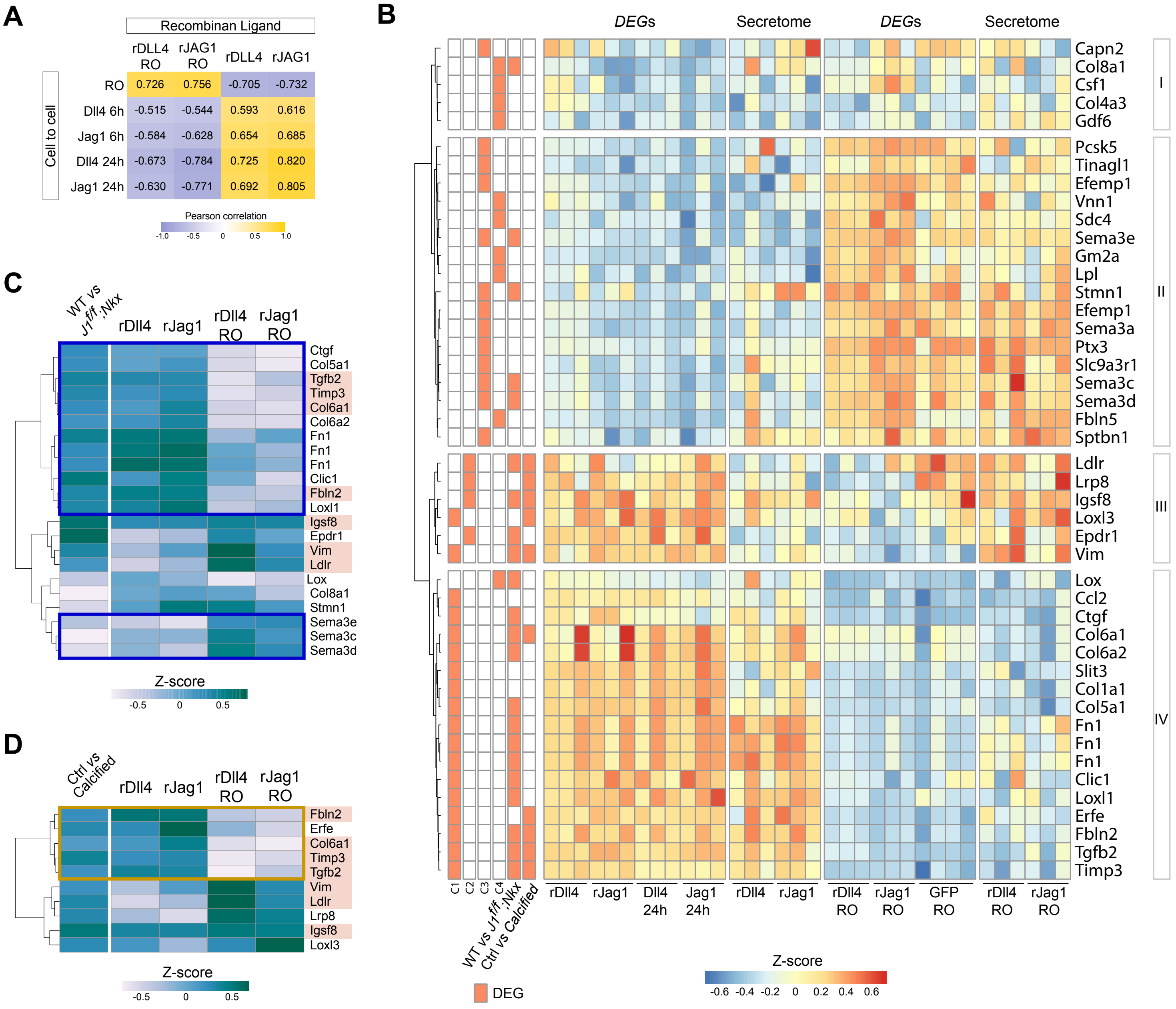


Luna-Zurita et al._Figure S2

**Figure S2. Notch dependent endocardial secretion and transcriptional changes**

(A) Pearson correlation between the changes in MEEC gene expression induced by co-culture with OP9-DLL4 and OP9-JAG1 cells for 6 and 24 hours.

(B) Hierarchical clustering of DEGs based on their changes in gene expression after Notch pathway stimulation or inhibition in the MEEC-OP9 co-culture system or with recombinant ligands and the differential secretion of the resultant proteins.

(C) Hierarchical clustering of C1-C4 DEGs differentially enriched in E14.5 WT or *Jag1$^{flox}$;Nkx2.5-Cre* semilunar valves showing differential Notch-dependent secretion.

(D) Hierarchical clustering of C1-C2 DEGs differentially enriched in Control versus calcified adult human valves showing differential Notch-dependent secretion.

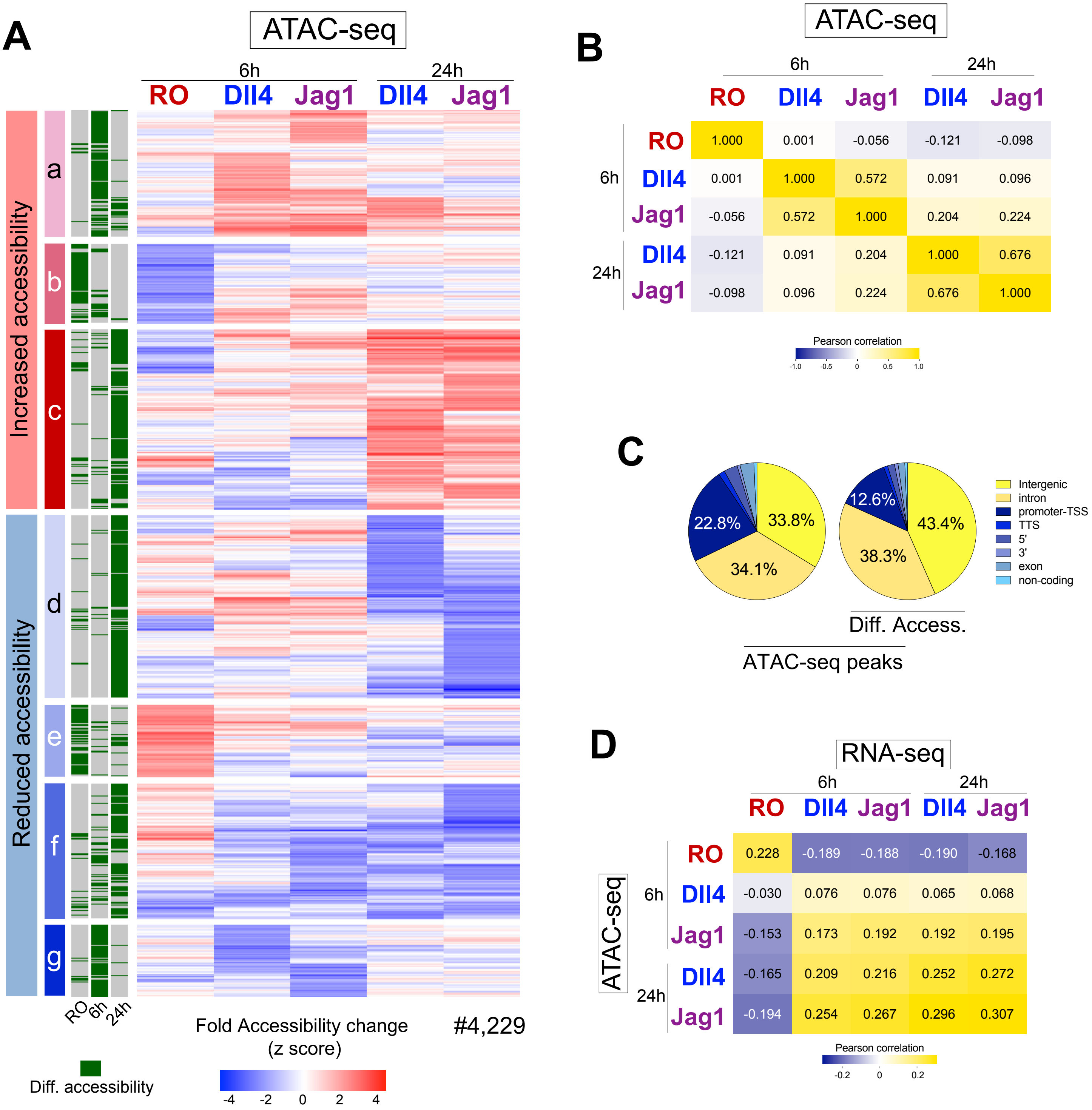


Luna-Zurita et al._Figure S3

**Figure S3. Chromatin accessibility changes in mouse embryonic endocardial cells after Notch-ligand activity**

(A) Hierarchical clustering of genomic regions showing differential accessibility in MEEC treated with RO or co-cultured with OP9 cells expressing Dll4 or Jag1.

(B) Pearson correlation between experimental conditions for changes in chromatin accessibility.

(C) Genome wide distribution of ATAC-seq peaks showing or not showing differential accessibility.

(D) Pearson correlation between the changes in chromatin accessibility in differential peaks and changes in gene expression of the associated DEGs in each experimental condition.

A

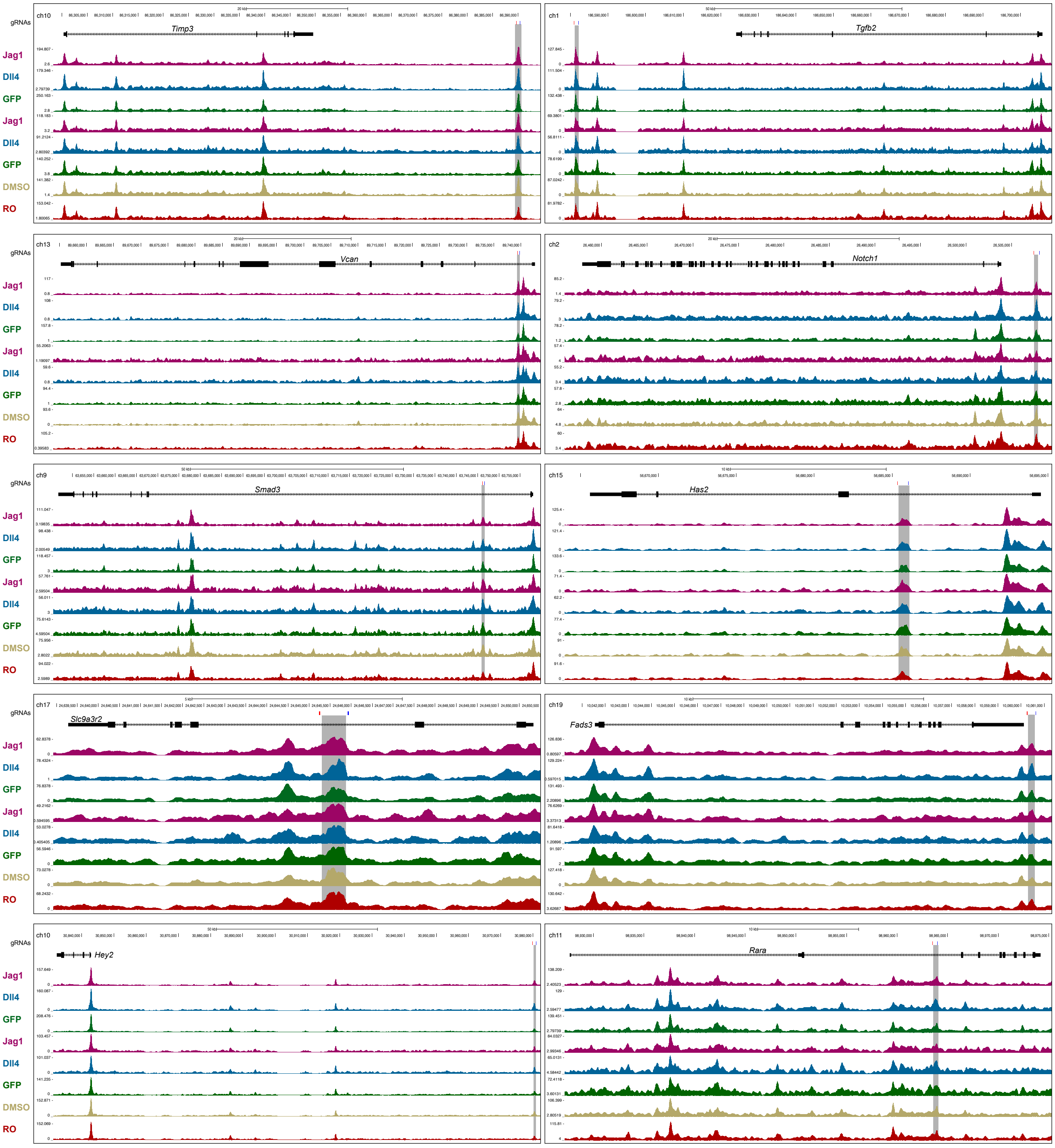


B

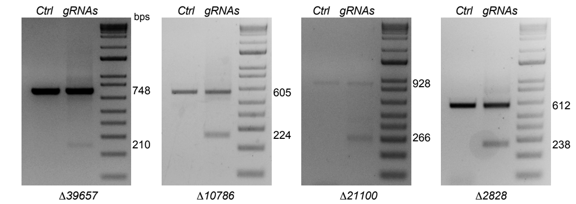


Luna-Zurita et al._Figure S4

**Figure S4. Transcriptional effect on target genes of regulatory-element deletion in mouse embryonic endocardial cells**

(A) Scheme in the mouse genome (mm10) indicating the location of the Notch-dependent deleted regulatory elements and of the guide RNAs.

(B) Representative PCR analysis of genomic DNA from MEEC transfected with empty pDG458 and pDG458 containing 2 specific guides for the region of interest.

**Supplementary Table Legends**

**Supplementary Table S1**. Gene expression analysis in MEEC-OP9 co-cultures
Sheet 1: DEGs in MEEC+OP9-Dll4 vs MEEC+OP9-GFP 6h co-cultures
Sheet 2: DEGs in MEEC+OP9-DJag1 vs MEEC+OP9-GFP 6h co-cultures
Sheet 3: DEGs in MEEC+OP9-Dll4 vs MEEC+OP9-GFP 24h co-cultures
Sheet 4: DEGs in MEEC+OP9-DJag1 vs MEEC+OP9-GFP 24h co-cultures
Sheet 5: DEGs in MEEC+OP9-GFP+RO vs MEEC+OP9-GFP+DMSO 6h co-cultures
Sheet 6: Combinations of DEGs detected in at least one condition. Log2(Norm.Counts) are presented
Sheet 7: GO-ANALYSIS cluster C1*
Sheet 8: GO-ANALYSIS cluster C2*
Sheet 9: GO-ANALYSIS cluster C3*
Sheet 10: GO-ANALYSIS cluster C4*
* Tables are the standard output from Metascape

**Supplementary Table S2**. In vivo analysis of DEGs detected in MEEC-OP9 co-cultures
Sheet 1: GSEA output from MEEC vs E14.5 mouse semilunar valves (WT *vs Jag1f/f;Nkx2-5-Cre*) and adult human valves (Control *vs* Calcified BAV and TAV)
Sheet 2: C1-C2 DEGs enriched in E14.5 mouse semilunar valves (WT *vs Jag1f/f;Nkx2-5-Cre*) and adult human valves (Control *vs* Calcified BAV and TAV)
Sheet 3: GO-ANALYSIS cluster A1*
Sheet 4: GO-ANALYSIS cluster A2*
Sheet 5: GO-ANALYSIS cluster A3*
Sheet 6: STRING database output (A1-A3)
* Tables are the standard output from Metascape

**Supplementary Table S3**. Secretome and RNA-seq in MEEC
Sheet 1: DEGs in MEEC+rJAG1 vs MEEC+DMSO
Sheet 2: DEGs in MEEC+rDLL4 vs MEEC+DMSO
Sheet 3: DEGs in MEEC+rJAG1 vs MEEC+rJAG1+RO
Sheet 4: DEGs in MEEC+rDLL4 vs MEEC+rDLL4+RO
Sheet 5: DEGs in MEEC+rJAG1+RO vs MEEC+DMSO

Sheet 6: DEGs in MEEC+rDLL4+RO vs MEEC+DMSO

Sheet 7: MEEC+OP9 24h (RNA-seq) – MEEC+rLIGANDS (RNA-seq) – MEEC+rLIGANDS (Secretome)

Sheet 8: E14.5 WT *vs Jag1f/f;Nkx2-5-Cre* (RNA-seq) – MEEC+rLIGANDS (Secretome)

Sheet 9: Adult Human valves Control *vs* Calcified BAV and TAV (RNA-seq) – MEEC+rLIGANDS (Secretome)

**Supplementary Table S4**. ATAC-seq in MEEC

Sheet 1: Differential Peaks in MEEC+OP9-GFP+RO vs MEEC+OP9-GFP+DMSO 6h

Sheet 2: Differential Peaks in MEEC+OP9-Dll4/Jag1 vs MEEC+OP9-GFP 6h

Sheet 3: Differential Peaks in MEEC+OP9-Dll4/Jag1 vs MEEC+OP9-GFP 24h

Sheet 4: Differential consensus peaks

Sheet 5: Known HOMER Motifs identified in each group of differential peaks

Sheet 6: Differential peaks-DEGs association by GREAT using default settings. Output from single nearest gene setting is indicated.

Sheet 7: Differential peaks-DEGs association by GREAT using default settings. Only Peaks from clusters a-c detected in vivo and DEGs from clusters C1-C2 represented in vivo are considered

**Supplementary Table S5**. Enhancer deletion in MEEC and zebrafish

Sheet 1: gRNAs, genotyping deletion primers, and qPCR primers
